# StarSignDNA: Signature tracing for accurate representation of mutational processes

**DOI:** 10.1101/2024.06.29.601345

**Authors:** Christian Domilongo Bope, Sumana Kalyanasundaram, Knut D. Rand, Sigve Nakken, Ole Christian Lingjærde, Eivind Hovig

**Affiliations:** Centre for Bioinformatics, University of Oslo, Oslo, Norway; Department of Informatics, Faculty of Mathematics and Natural Sciences, University of Oslo, Norway; Department of Tumor Biology, Institute for Cancer Research, The Norwegian Radium Hospital, Oslo University Hospital, Oslo, Norway; Centre for Cancer Cell Reprogramming, Institute of Clinical Medicine, Faculty of Medicine, University of Oslo, Oslo, Norway; Department of Mathematics and Department of Computer Science, Faculty of Sciences, University of Kinshasa, Kinshasa, Democratic Republic of Congo; Department of Cancer Genetics, Institute for Cancer Research, Oslo University Hospital, Oslo, Norway

## Abstract

All cell lineages accumulate mutations over time, increasing the probability that some lineages eventually become malignant. Many of the processes responsible for generating mutations are leaving a characteristic footprint in the genome that allows their presence to be detected. However, the mutational pattern in a tumour is usually the combined result of multiple mutational processes being at work simultaneously, and the problem of disentangling the different footprints and their relative impact then becomes a deconvolution problem. Several algorithms have been developed for this purpose, most of them involving a factorization of the mutation count matrix into two non-negative matrices, representing respectively the underlying mutational signatures and the relative weighting of (or exposure to) these signatures. Here, we introduce the StarSignDNA algorithm for mutational signature analysis, which offers efficient re-fitting and *de novo* mutational signature extraction. StarSignDNA is capable of deciphering well-differentiated signatures linked to known mutagenic mechanisms and suggesting clinically relevant predictions for a single patient. The package offers a command line-based interface and data visualization routines.

**Author summary:** StarSignDNA is a novel algorithm for identifying mutational signatures from cancer sequencing data. It excels in low mutation count scenarios, balancing signature detection and true signature discovery. StarSignDNA improves prediction accuracy and biological validity by addressing challenges such as overfitting and underfitting, achieving optimal variable selection and shrinkage, and ensuring interpretability. In re-fitting, it reduces prediction variance and handles sparsity, resulting in more precise predictions. In *de novo* extraction, it improves the detection of challenging signatures and achieves better alignment between detected and known signatures through unsupervised cross-validation. Its unique features, including prediction confidence and customizable reference signatures, offer valuable insights for clinical single-sample analysis.

## Introduction

A diverse set of mutational processes shape cells that ultimately may develop into cancers. These processes encompass endogenous cellular mechanisms, such as errors in the DNA replication and repair machinery, as well as exogenous sources of DNA damage, like tobacco and UV exposure [1]. The imprints of mutational processes that have occurred during tumourigenesis are reflected in distinct spectra of acquired DNA aberrations in tumour samples, and these aberrations are now effectively captured with high-throughput sequencing technologies. A characteristic pattern of mutations caused by a mutational process is now commonly referred to as a mutational signature [2]. Accurate identification of mutational signatures across tumour samples is critical for our understanding of the mechanisms driving tumour development, and it can also add novel value when it comes to patient stratification toward personalized treatment regimes [3]. While distinct patterns caused by somatic copy number aberrations and insertion/deletion type of mutations have been found [4], we here focus on the signatures exerted by single base substitutions (SBS), the most common type of aberration in cancer genomes.

Several algorithms and computational tools have been developed to detect and estimate mutational signatures in cancer genomes. These methods can broadly be classified into two main categories: discovery of novel signatures in a collection of sequenced tumour samples (also referred to as *de novo* signature identification), and fitting known reference signatures to an existing mutational profile (signature reconstruction or re-fitting). Non-negative Matrix Factorization (NMF), or its Bayesian version (BNMF) [1], [5–7] [8–13], is a matrix decomposition approach that breaks down a complex matrix into smaller, more manageable subproblems. Apart from NMF-based approaches, probabilistic methods such as *Emu*, use Expectation Maximization. In *Emu*, the underlying probabilistic model assumes that input samples are independent and that the number of mutational signatures is inferred using the Bayesian Information Criterion (BIC) [14]. Another method, *pmsignature*, employs a flexible approach that simultaneously reduces the number of estimated parameters and allows for the modification of key contextual parameters, such as the number of flanking bases [15]. The collective effort of identifying signatures is collated in the Catalogue Of Somatic Mutations In Cancer (COSMIC, https://cancer.sanger.ac.uk). COSMIC also links each signature to the proposed underlying processes. The re-fitting algorithms aim to search for the best combination of established COSMIC signatures that explains the observed mutational catalogue.

However, despite advances in the field, challenges remain in accurately characterizing mutational signatures. A key challenge is to robustly recover the signature compositions when the mutation data is limited, which is a common scenario for many tumor samples sequenced through whole-exome or targeted assays. A low mutation count can lead to unreliable discovery of some active signatures, increased prediction variance, and higher fitting error [16]. To address this challenge, a common practice in signature discovery is the use of regularisation. Regularisation promotes sparsity in the model and essentially encourages fewer non-zero coefficients. This can help to prevent overfitting, but it can also introduce a bias into the predictions which can be negligible when using a better optimization approach. Several methods achieve regularisation in different ways; for example, by introducing constraints on the model, by using a Bayesian variant of Non-Negative Matrix Factorization (NMF) with a penalty on model complexity, by employing an L1-based regularisation for sparsity on the signatures, or by leveraging the L1 norm penalty to promote sparsity [12], [17], [18]. These techniques aim to achieve better predictions by reducing the estimates’ variance.

In addition to the aforementioned challenges, current *de novo* signature discovery methods face the following weaknesses: (i) Determining the optimal number of signatures (K) to extract; there is currently no definitive method for the selection of K, (ii) Balancing the number of signatures (K) and their similarity; achieving a good balance between K and their similarities to known signatures (measured by cosine similarity, typically with a relatively high threshold (e.g. 0.8) is desirable. It is important to remember that simply fitting the data well (minimizing residual error) doesn’t necessarily reflect actual biological processes [11] [19]. The common practices to determine the number of signatures are: (1) The number of signatures (K) should be selected in such a way that higher K will imply lower residual error [11]; (2) K is selected based on the residual error and maximizing the reproducibility of signatures; (3) Calling signatures hierarchically on subsets of samples, adding more signatures to fit every sample. The first two approaches are arbitrary and the third leads to overfitting, since there is a need to use more signatures for better fitting without any constraint. Recently, different methods have been developed to improve the selection of the number of signatures, namely SignatureAnalyzer [5] and SUITOR [20].

SignatureAnalyzer uses automatic relevance determination, which starts with a high number of signatures and attempts to eliminate signatures of low relevance [5]. SparseSignatures employs cross-validation to jointly determine the optimal number of signatures and shrinkage parameter. This approach, while computationally demanding, often leads to the identification of signatures with pronounced peaks [12]. SUITOR is an unsupervised cross-validation method that selects the optimal number of signatures to attain the minimal prediction error in the validation set and extends the probabilistic model to allow missing data in the training set.

In this paper, we introduce StarSignDNA, an NMF model that offers *de novo* mutation signature extraction, as well as re-fitting to existing mutational signatures. The algorithm uses regularization for its dual benefits: variable selection and shrinkage [17, 21, 22]. Variable selection is essential in mutational signature analysis for identifying the most relevant signatures among many possible ones, enhancing interpretability. Shrinkage helps prevent overfitting by reducing the influence of weak signatures. StarSignDNA combines the use of L1-regularisation with the Poisson model to allow stable estimates for the data to accommodate low mutational counts.

StarSignDNA overcomes the aforementioned limitations for both re-fitting and *de novo*. StarSignDNA re-fitting is optimized in the following ways:

1. Utilizing LASSO regularization to minimize the spread (variance) in exposure estimates. This helps ensure more consistent and reliable exposure values, reducing the overall variability.
2. Stable exposure estimation. This ensures a minimum number of accurate processes with low fitting errors to describe the mutational catalogue of a cancer genome.
3. Improved F1 score by reducing both false positives (overfitting of signatures) and false negatives.
4. Single-Patient Suitability which provides confidence levels on the predicted processes, making it suitable for single-patient evaluation of mutational signatures.
5. Weight matrix integration is the incorporation of the distribution of triplets in a reference genome/exome or normal tissue from the same patient (weight matrix).

The StarSignDNA *de novo* module is optimized in the following ways:

1. Use the optimal model selection by combining unsupervised cross-validation and the probability mass function as a loss function to select the best combination of the number of signatures (K) and regularisation parameters (*λ*).
2. The algorithm avoids introducing bias towards unknown signatures and achieves a better balance between the number of signatures (K) and the cosine similarity threshold (above 0.8).
3. Demonstrates high sensitivity in detecting signatures.

The package is available at https://github.com/uio-bmi/StarSignDNA and can be installed through PyPI.

## Methods and materials

### Mutational signatures

Single base mutations can be classified in 96 categories *X*[*Y* → *Z*]*W*, where Y denotes the reference nucleotide, Z is the mutated variant, and X and W represent the two flanking nucleotides. Let **m** = (*m*_1_, …, *m*_96_) ∈ ℝ^96^ denote the number of mutations in each category, with the convention that the categories are lexicographically sorted on the mutation names. Suppose this mutation pattern is the combined effect of several mutational processes with distinct mutational signatures **s**_*i*_, *i* = 1, …, *K*. Then the observed mutational spectrum **m** may be modeled as a random sample from a distribution *p*(**m**|***θ***) with mean ***µ*** = *e*_1_**s**_1_ + … + *e*_*K*_**s**_*K*_, where all the weights *e*_*i*_ are non-negative and some may be equal to zero. Both the signatures and the weights may require estimation from a collection of observed mutational patterns (*de novo* signature extraction). If the signatures are known, we only have to estimate the weights (signature re-fitting). The proposed method can perform both operations.

Distinct mutagenic processes are assumed to leave different footprints in the genome in the form of different mutational signatures. Each mutational signature is a numerical vector **s**_*k*_ = (*s*_1*k*_, …, *s*_96,*k*_)^*T*^ ∈ ℝ^96^, specifying the relative frequency of occurrence of each of the 96 mutation types. With no loss of generality, we may assume that the components of a signature sum to 1 (this implicitly requires that at least one mutation type is present in a signature, which is a reasonable assumption).

Suppose we have a vector **m** = (*m*_1_, …, *m*_96_)^*T*^ of mutation counts for a single sample and also a 96 *× K* signature matrix **S** = (*s*_*jk*_). We wish to determine the exposure vector **e** = (*e*_1_, …, *e*_*K*_)^*T*^ where each element is the non-negative weight given to a particular signature. If we assume that the mutation counts follow a normal distribution *p*(**m**|***θ***) = 𝒩 (**m**|***µ*, Σ**), with a sample-specific mean ***µ*** and the variance-covariance matrix **Σ** = *σ*^2^**I**, where **I** is the identity matrix, the maximum likelihood estimate of the exposure vector is the minimizer of

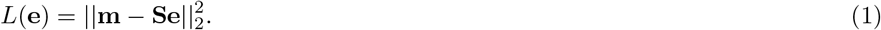

In practice, the normality assumption depends on the counts being sufficiently separated from zero. Mutational data obtained from low-coverage sequencing may not satisfy this assumption. To circumvent this problem, other distributional assumptions may be made. For example, *sigfit* [23] assumes that the observed mutation counts follow either a Gaussian, a Poisson, or a negative binomial distribution, while Li et al. [17] assumes that the mutations follow a multinomial distribution.

An early example of regularization in NMF is the method Sparse Non-negative Matrix Factorization *SNMF/L* [18]. For known signature matrix **S** this method applies the loss function

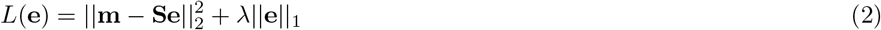

The penalty term imposes a soft constraint on the *L*^1^-norm of the exposure vector **e**. The effect of this constraint is to force the solution to balance a good fit to the data (the first term) with a small norm of **e**. The *L*^1^-norm will tend to force several components of **e** to be zero. Thus, the model will produce a sparse NMF decomposition in **e**. In the closely related method *SNMF/R* the loss function is

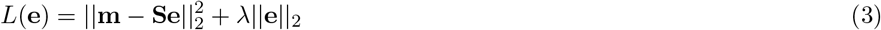

which results in an NMF decomposition that is not sparse in **e**. Another method is *sigLASSO* [17] which assumes a multinomial distribution of the mutation counts and with a Laplacian prior on the exposure levels.

### Estimating the exposure matrix E

We first consider the problem of determining the optimal exposure matrix **E**, given a signature matrix **S** and mutation counts for a set of samples. The optimal exposures for a sample will not depend on the mutation counts for other samples, and the problem can be solved for a single sample and then extended trivially to handle multiple samples by iterating through samples.Suppose, then, that we have observed mutation counts *m*_1_, …, *m*_96_ for a given sample. These are assumed to follow Poisson distributions Poi(*m*_*j*_|*µ*_*j*_) with unknown rates *µ*_*j*_ satisfying the system of equations

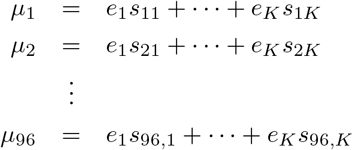

Here, *s*_*jk*_ are known signature values (assumed to be non-negative), and *e*_*k*_ ≥ 0 are unknown sample-specific parameters. To ensure the uniqueness of the solution, we must require *K* ≦ 96, i.e., the number of signatures should be no larger than the number of mutation categories. Assuming that the signature matrix *S* = (*s*_*jk*_) has rank *K* we may find the *e*_*k*_ by maximizing the log-likelihood

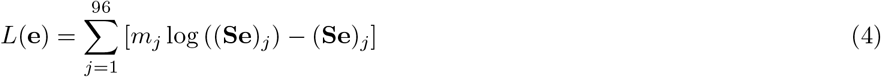

over 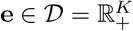.

To find a sparse non-negative solution, we could instead maximize the *L*^1^-penalized log-likelihood

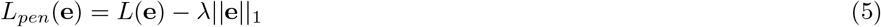

over **e** ∈ 𝒟, where *λ >* 0 controls the trade-off between goodness-of-fit and sparsity. This function is differentiable on 𝒩 = 𝒟 *\* {**e** : *e*_*k*_ = 0 for at least one *k*}. By a suitable extension of the gradient **g**(**e**) on 𝒩 to a function 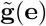 defined on the whole region 𝒟, we may solve the constrained penalized optimization problem in Eq 5 by gradient ascent, i.e. by updates of the form 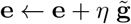.

We define 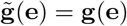 whenever **e** ∈ 𝒩 and define 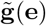 to point into the feasible region for border points **e** ∈ {**e** : *e*_*k*_ = 0 for at least one *k*}. Specifically, we first compute the gradient **h**(**e**) = (*h*_1_, …, *h*_*K*_)^*T*^ of the unpenalized likelihood:

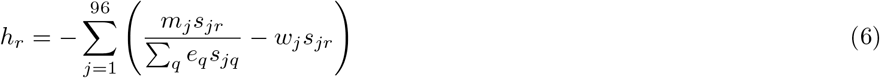

and second, we calculate the extended gradient of the penalized likelihood as 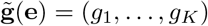 where

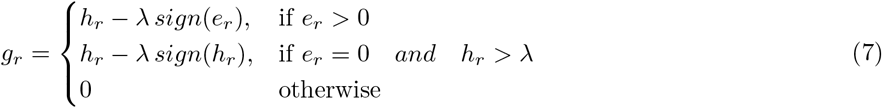

where *sign* (*x*) is -1, 0, or 1 depending on whether the argument *x* is negative, zero, or positive. Consider first the case where the current solution **e** is in the interior of the feasible region. Then **e** ∈ 𝒩 and *e*_*r*_ *>* 0 for all *r*, hence *g*_*r*_ = *h*_*r*_ − *λ* and 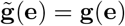. In this case, we see that the extended gradient coincides with the gradient of the penalized likelihood. If, on the other hand, the current solution is on the border of the feasible region, we have *e*_*r*_ = 0 for at least one *r*. If the gradient of the unpenalized likelihood is large and points into the feasible region (more specifically, *h*_*r*_ *> λ*), then the extended gradient of the penalized likelihood is *g*_*r*_ = *h*_*r*_ − *λ >* 0 and points into the feasible region. If the gradient of the unpenalized likelihood is small and pointing into the feasible region (specifically, 0 ≤ *h*_*r*_ ≤ *λ*) or pointing out of the feasible region (*h*_*r*_ *<* 0), we have *g*_*r*_ = 0 implying that the next gradient ascent iteration will not change the *r*th component of the solution vector. To ensure that each iteration of the gradient ascent algorithm results in a novel feasible solution, we must ensure that each component *e*_*r*_ of the solution remains non-negative after the update *e*_*r*_ ← *e*_*r*_ + *η g*_*r*_. Hence, we must ensure that *η g*_*r*_ *>* −*e*_*r*_, or equivalently that *η >* −*e*_*r*_*/g*_*r*_, for all *r*. This is accomplished by bounding the learning rate *η* to not exceed

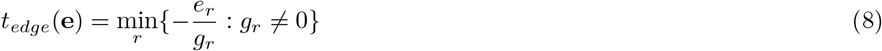

where *g*_*r*_ is the *r*th component of **g**(**e**). To ensure that the updated solution is close to the optimum, we also consider the curvature of the penalized likelihood at the current solution along the extended gradient direction, essentially choosing the learning rate to match what the optimal learning rate would be if the cost function was quadratic:

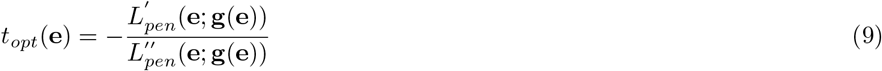

The actual learning rate is chosen to be *η* = min(*t*_*edge*_, *t*_*opt*_).

The optimisation starts with an initial value **e** = **e**^(0)^ for the solution. Each iteration of the optimization takes one of two forms. If the current solution is well separated from all the borders *e*_*k*_ = 0, *k* = 0, 1, …, *K*, then a gradient ascent step is performed:

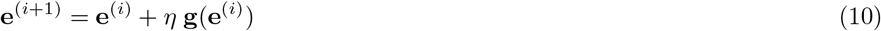

where *η >* 0 is the learning rate. If, on the other hand, the current solution is close to a border, a Newton-Raphson step is performed:

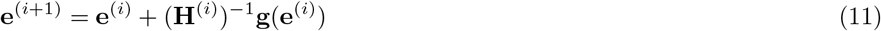

where **H**^(*i*)^ is a modified version of the Hessian of the penalized likelihood calculated in **e**^(*i*)^. The modification entails replacing second partial derivatives 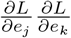 in **H**^(*i*)^ with 0 whenever 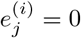 or 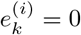, hence leaving the current solution at rest in dimensions where the solution is on the border.

### Estimating the signature matrix S

We next consider the problem of estimating the optimal signature matrix **S** for a given exposure matrix **E**. We start by reformulating the optimization problem considered in the previous section. Suppose we have mutation counts *m*_*ij*_ for samples *i* = 1, …, *n* and mutation categories *j* = 1, …, 96. According to Eq 4, the optimal exposure levels **e**_*i*_ can be found by maximizing the expressions

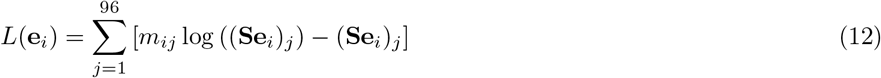

for *i* = 1, …, *n*, or equivalently to maximize

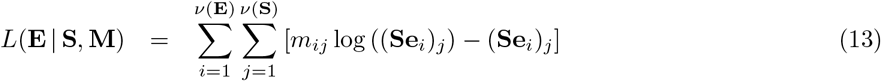

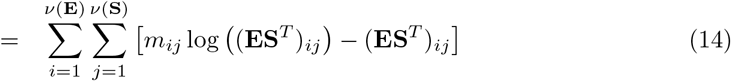

where *ν*(**E**) and *ν*(**S**) denotes the number of rows in the matrices **E** and **S**, respectively. In the previous section, we described how to determine the optimal **E** in this expression for given **S** and **M**. In particular, we may apply this method to determine the optimal solution of

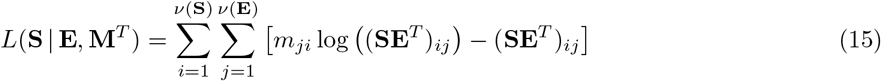

Switching the names of the two indices *i* and *j* and the order of the two sums, we obtain the equivalent expression

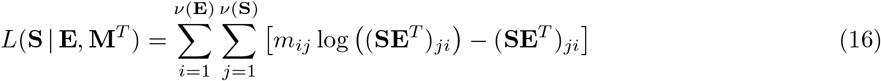

which is easily seen to be identical to *L*(**E** | **S, M**). Accordingly, maximizing *L*(**S** | **E, M**^*T*^) with respect to **S** is equivalent to maximizing the original likelihood expression Eq 13 with respect to **S**. In conclusion, we may estimate the signature matrix **S** using the method already described to estimate the exposure matrix **E**.

### StarSignDNA algorithms

StarSignDNA consists of four main algorithms designed for different analytical tasks. Algorithm 1 is used for single-sample re-fitting, allowing the model to be adjusted using data from a single sample. Algorithm 2 extends this capability to multiple samples, enabling multi-sample re-fitting, which simultaneously refits the model across several datasets. Algorithm 3 focuses on discovering new patterns by performing *de novo* signature estimation across multiple samples, which helps identify novel signatures. Lastly, Algorithm 4 is dedicated to optimizing the model by estimating the best values for the number of signatures (*K*) and the regularization parameter (*λ*), ensuring that the model is both accurate and efficient.

#### Algorithm 1

Single-sample re-fitting

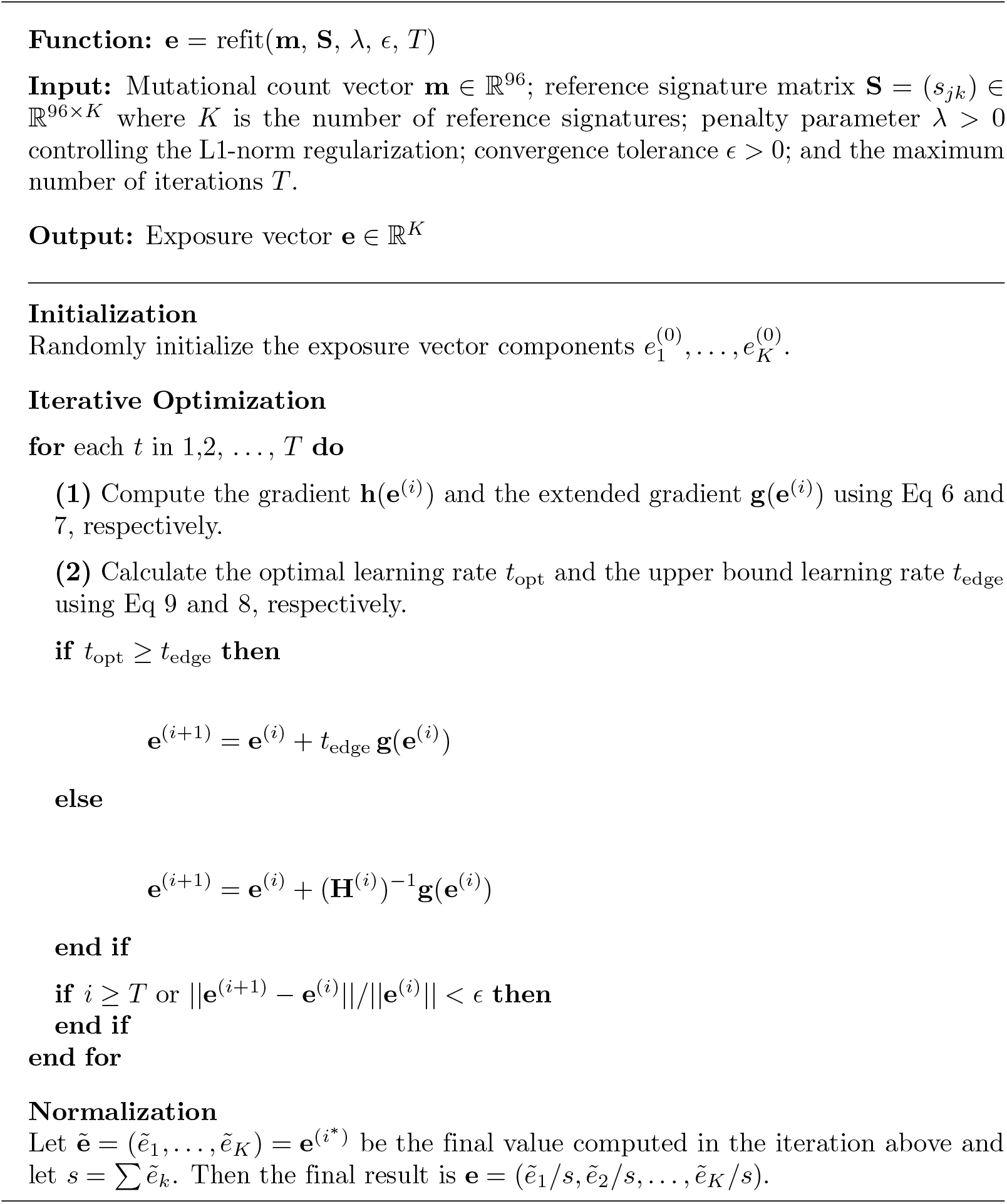

#### Algorithm 2

Multi-sample re-fitting

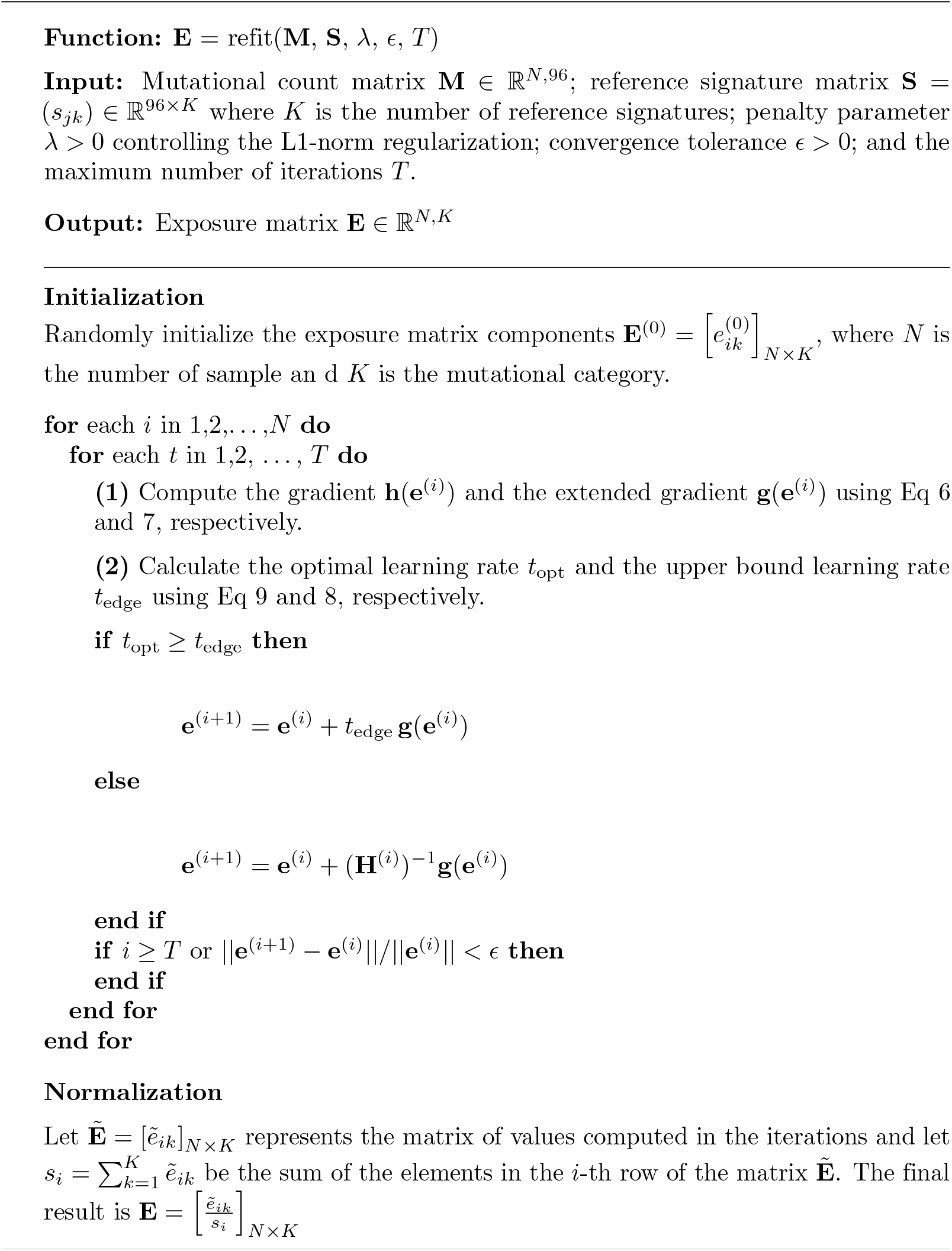

#### Algorithm 3

*De novo* Mutational Signatures with fixed *K* and *λ*

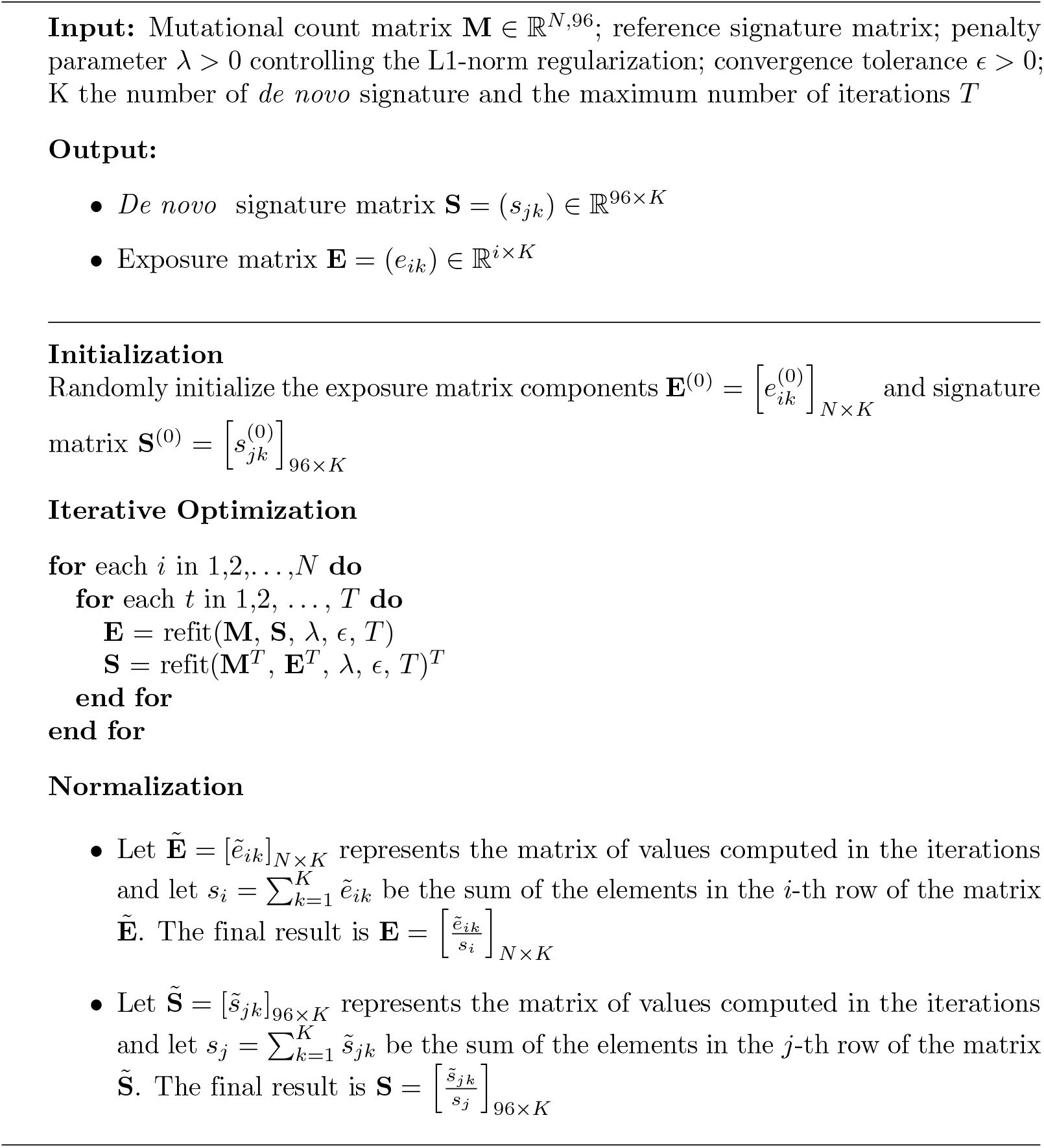

#### Algorithm 4

Estimation of *K* and *λ*

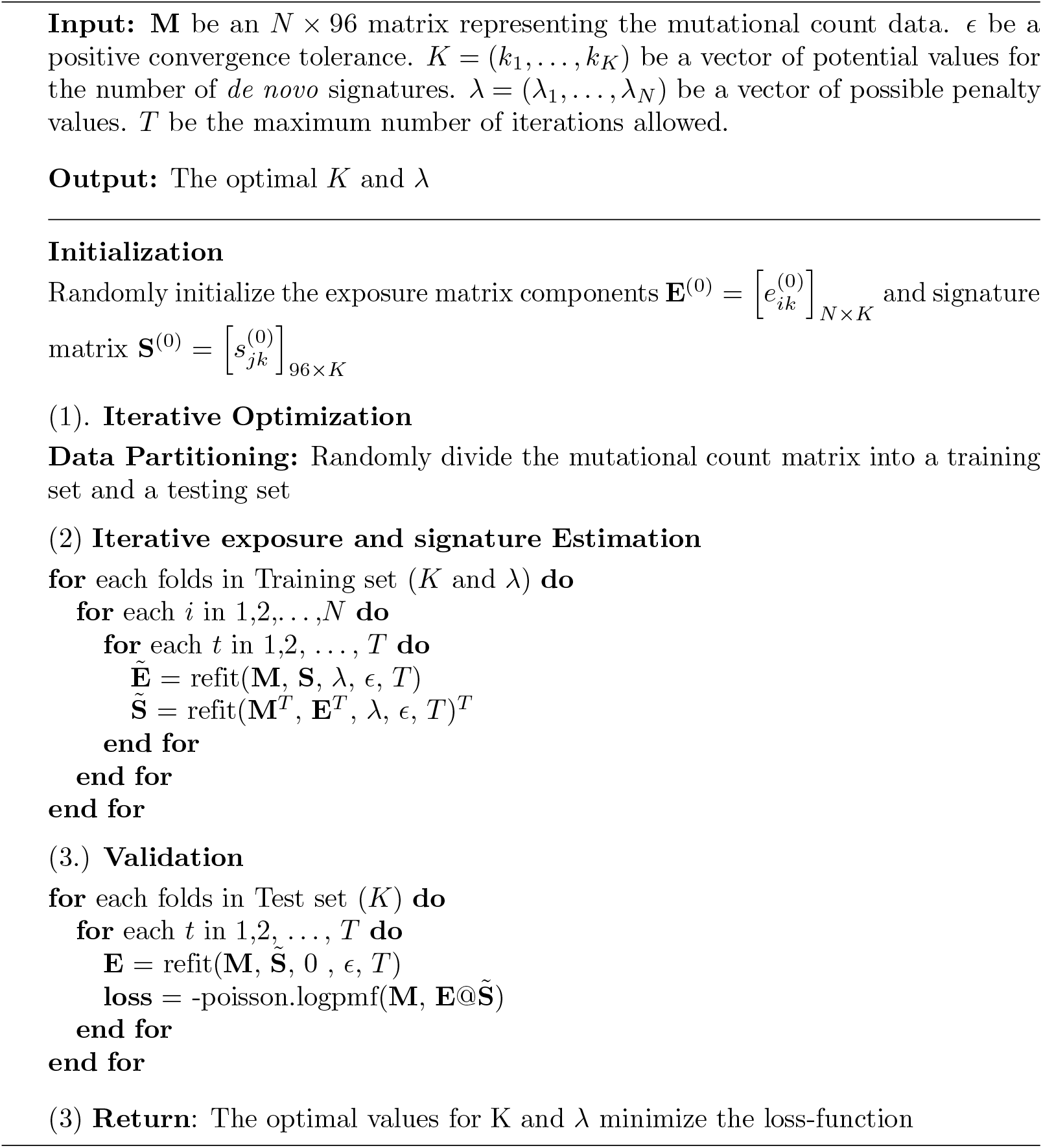

## Materials

StarSignDNA’s performance was assessed using both simulated data and publicly available mutation datasets from tumor samples.

### Simulated data

We first evaluated the performance of StarSignDNA using multiple simulation datasets. For a fair and consistent comparison of different methods, it is crucial to have true underlying processes as a reference and then evaluate the ability of the various methods to detect them. Five categories of simulated datasets were considered. The first four categories were used to validate the re-fitting algorithm, and the last was used to assess the *de novo* algorithm. To generate the datasets for the first three scenarios, we employed *N* mutation vectors for each three-mer context. Here, *N* represents the total number of mutations observed in that specific context. Each component of the *N*^*th*^ mutation vector was drawn from a non-negative binomial distribution with *N* trials and a success probability determined by the corresponding component of the *N*^*th*^ mutation vector. We introduced a vector of exposure values, **e**_*_ = (*e*_1_, *e*_2_, …, *e*_*R*_), and calculated the mean, *µ*, as a linear combination of the selected reference signatures (SBS) and these exposure values: *µ* = *e*_1_ · *SBS*1 + *e*_2_ · *SBS*2 + … + *e*_*R*_ · *SBS*_*R*_. The variance for each signature was set to 10.1 matching the observed variance in real-world mutation data.

The fourth category of simulations was generated using the recently developed GENOMICON-Seq [24], a mutation simulator tool that incorporates laboratory protocol noise. The noise encompasses probe-capturing enrichment bias (only fragments with a 100% match to the probes were selected), indexing PCR amplification bias over 8 cycles, sequencing bias where shorter fragments have a higher probability to be sequenced, and sequencing error mimicking the Illumina NovaSeq instrument.

The first category consisted of 16 datasets (Table 1), which incorporated variations in exposure weights and the number of mutations per sample (K=100, 500, 1000, and 10000). As mentioned earlier, K is the number of mutations per sample to analyze the sensitivity and precision.

**Table 1.** The first category of simulated mutation datasets, consisting of 16 datasets, each containing 100 samples. These datasets were drawn from a negative binomial distribution and were used to explore the impact of two factors: exposure values and the number of mutations per sample. Each row represents a different set of exposure values, while each column represents one of the four mutational signatures. The number of mutations (K=100).

| Dataset | SBS1 | SBS2 | SBS3 | SBS4 | Mutations (K) |
| --- | --- | --- | --- | --- | --- |
| SimA | 1 | 0 | 0 | 0 | 100 |
| SimB | $\frac{1}{2}$ | $\frac{1}{2}$ | 0 | 0 | 100 |
| SimC | $\frac{1}{3}$ | $\frac{1}{3}$ | $\frac{1}{3}$ | 0 | 100 |
| SimD | $\frac{1}{4}$ | $\frac{1}{4}$ | $\frac{1}{4}$ | $\frac{1}{4}$ | 100 |

In the second (Table 2) and third categories (Table 3), we considered two subsets of the COSMIC reference signatures version 3.4 with different characteristics (S1 Figure). The first subset consisted of signatures close to all other signatures (hence challenging to uniquely identify), while the second subset consisted of signatures distinct from all other signatures, and thus less challenging to detect. Technically, the first subset was found by identifying the five signatures with the highest average cosine similarity to all other signatures and the highest cosine similarity to at least one other signature. In comparison, the second subset was found by identifying the five signatures with the lowest average and maximum cosine similarity to the different signatures. The first subset consists of the signatures SBS3, SBS5, SBS40c, SBS92, and SBS94, which all have broad support (many non-zero components). Such signatures are commonly referred to as flat signatures. The second subset consists of the signatures SBS7d, SBS10b, SBS13, SBS17b, and SBS28, which all have narrow support (few non-zero components). Such signatures are here referred to as common signatures since they are more easily identifiable.

**Table 2.**
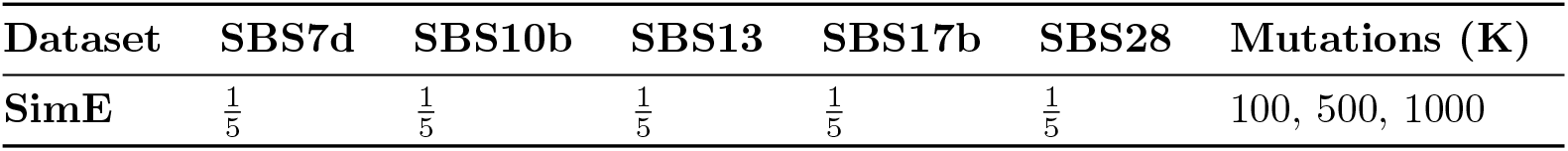
The second category of simulated mutation datasets, consisting of 9 datasets, each containing 100 samples. The row represents exposure values, while each column represents one of the five mutational signatures. The number of mutations (K) varies within these datasets.

| Dataset | SBS7d | SBS10b | SBS13 | SBS17b | SBS28 | Mutations (K) |
| --- | --- | --- | --- | --- | --- | --- |
| SimE | $\frac{1}{5}$ | $\frac{1}{5}$ | $\frac{1}{5}$ | $\frac{1}{5}$ | $\frac{1}{5}$ | 100, 500, 1000 |

**Table 3.** Performance using the third category of simulated mutation datasets, consisting of seven datasets, each containing 100 samples. The row represents exposure values, while each column represents one of the five mutational signatures. An additional layer of complexity is introduced by varying the total number of mutations within each dataset. The number of mutations (K) varies within these datasets. For instance, K = 50 indicates datasets where only 50% of the mutations from a baseline scenario (K = 100) are retained. Similarly, K = 70, K = 80, and K = 90 represent datasets with 70%, 80%, and 90% of the mutations remaining compared to the baseline, respectively.

| Dataset | SBS3 | SBS5 | SBS40c | SBS92 | SBS94 | Mutations (K) |
| --- | --- | --- | --- | --- | --- | --- |
| SimE | $\frac{1}{5}$ | $\frac{1}{5}$ | $\frac{1}{5}$ | $\frac{1}{5}$ | $\frac{1}{5}$ | 50,70,80,90,100, 500, 1000 |

The fourth category of simulated data generated samples containing 1,000 copies of the hg38 genome, with 80% of the genome harboring at least one mutation. To mimic SBS17b (easy signature to locate) and SBS92 (hard signature identify) mutational signatures, 3% of the total exonic positions were altered. All samples had an average sequencing depth of 30x. The *in-silico* FASTQ files were aligned using BWA [25], and variants were called using Strelka2 [26] by comparing the mutated (tumour) samples to mutation-free (control) samples.

The fifth category of simulated data was used to evaluate and compare the performance of *de novo* signature detection. For this, we considered *N* = 146 prostate samples, subject to whole-genome sequencing (WGS) within the PCAWG. We then selected a set of four signatures, namely SBS1, SBS5, SBS18, and SBS40, from the COSMIC signatures known to be prevalent in prostate cancer [5] (see supporting document). To ensure consistent comparison, 50 replicates were considered [16].

### Mutational signatures in patient-derived exomes and genomes

In total, 8,893 whole-exome sequencing (WES) samples across 33 cancer types from The Cancer Genome Atlas (TCGA) (https://portal.gdc.cancer.gov/) were used to evaluate the re-fitting method. A single TCGA breast cancer sample was used to demonstrate StarSignDNA’s ability to incorporate confidence into predictions using the 2.5% percentile to the 97.5% percentile of the bootstrapped exposure levels.

The *de novo* algorithm of StarSignDNA was evaluated using 3,308 whole-genome sequenced tumor samples selected from The Pan-Cancer Analysis of Whole-Genomes (PCAWG) and the Hartwig Medical Foundation (HMF) (Table 5).

**Table 4.** Comparison of methods on simulated datasets. The table presents the median per-patient correlation measured the correlation coefficient between the observed mutation spectrum and the predicted mutation spectrum for each patient.

| Methods | SimF(100) | SimF(500) | SimF(1000) |
| --- | --- | --- | --- |
| StarSignDNA | 0.51 | 0.76 | 0.87 |
| sigLASSO | 0.00 | 0.69 | 0.81 |
| deconstructSigs | 0.41 | 0.75 | 0.86 |
| MutationalPatterns | 0.41 | 0.75 | 0.86 |
| Sigfit | 0.40 | 0.72 | 0.83 |
| SigProfilerSS | 0.43 | 0.74 | 0.83 |

**Table 5.** WGS data used to validate the *de novo* algorithm. The HMF and PCAWG datasets include cancer types such as breast cancer (BRCA), kidney renal clear cell carcinoma (KIRC), pancreatic adenocarcinoma (PAAD), prostate adenocarcinoma (PRAD), and skin cutaneous melanoma (SKCM).

| Source | Type | BRCA | KIRC | PAAD | PRAD | SKCM |
| --- | --- | --- | --- | --- | --- | --- |
| HMF | WGS | 792 | 612 | 186 | 402 | 327 |
| PCAWG | WGS | 214 | 145 | 237 | 286 | 107 |

## Results

### Re-fitting Mutational signatures

#### StarSignDNA validation on simulated data

Considering the first category of simulated datasets (Table 1), which included variable exposure weights and number of mutations per sample, we evaluated the precision and sensitivity of six commonly used algorithms for mutational signature re-fitting: StarSignDNA, deconstructSigs, MutationalPatterns, sigLASSO, SigProfilerSingleSample (SigProfilerSS), and sigfit.

The weights assigned to the four signatures (SBS1, SBS2, SBS3, and SBS4) represent the ground truth. We employed a wider range of weights encompassing scenarios with high sensitivity and signature profile deviations to achieve a more comprehensive comparison. Furthermore, we investigated the impact of the number of mutations per sample on the performance of the different algorithms, considering scenarios with 100, 500, 1000, and 10000 mutations per sample. The results of the investigation are shown in (Fig 1) where we observe all the methods perform well, except SigProfilerSS and sigLASSO (Fig 1C and Fig 1D), which show some deviation from the ground truth. These results indicate that StarSignDNA, MutationalPatterns, sigfit, and deconstructSigs outperform SigProfilerSS and sigLASSO when more than two signatures are considered. When considering the higher number of mutations (K=500, 1000 and 10000) the result is shown in (S1 Figure, S2 Figure and S3 Figure), the performance is improved for SigProfilerSS and sigLASSO, indicate the challenges in both methods when a sample has a lower number of mutations (≈ 100).

**Fig 1.**
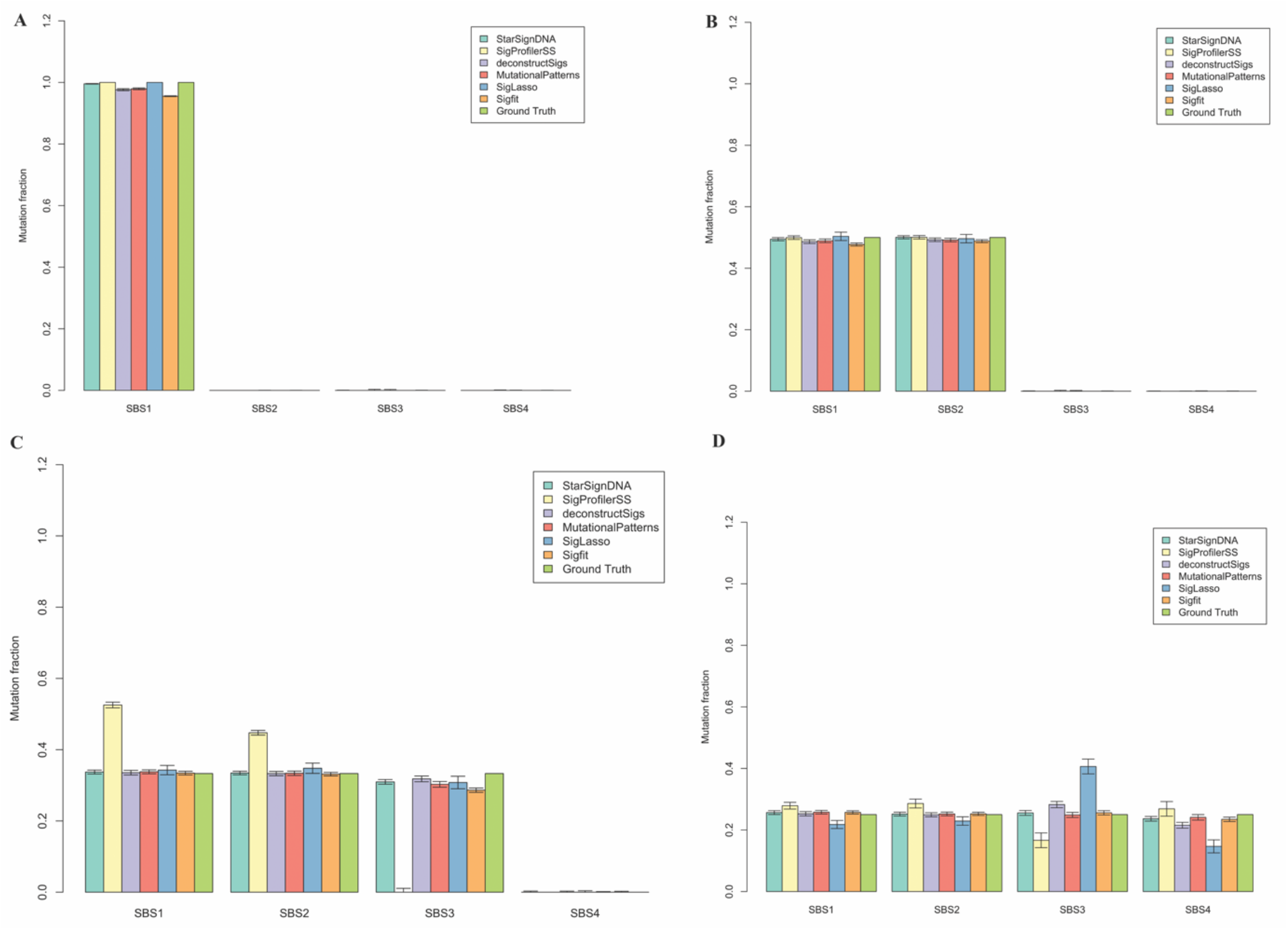
Performance comparison of six mutational signature re-fitting tools. StarSignDNA (light green), SigProfilerSS (light yellow), deconstructSigs (light blue), MutationalPatterns (red), SigLASSO (dark blue), and sigfit (orange) across four different signature types (SBS1, SBS2, SBS3, and SBS4) is represented by each set of bars. The vertical lines on top of each bar represent the average exposure of 100 simulated samples (approximately 100 mutations each) indicating the variability or uncertainty in the mutation fraction estimation by each tool compared to the known exposure values (Ground Truth). A) Only SBS1 has a true exposure of 1, while SBS2, SBS3, and SBS4 have a true exposure of 0. B) SBS1 and SBS2 have true exposures of 0.5, while SBS3 and SBS4 have true exposures of 0. C) SBS1, SBS2, and SBS3 each have true exposures of 0.333, while SBS4 has a true exposure of 0. D) All four signatures have an equal true exposure of 0.25.

We assess the performance of re-fitting algorithms on common signatures namely SBS7d, SBS10b, SBS13, SBS17b, and SBS28 more easily identifiable. We employed the second category of simulated data for this analysis (Table 2). The results are illustrated in Fig 2A-C which show StarSignDNA, deconstructSigs, MutationalPatterns, and sigfit achieved better prediction while SigProfilerSS and sigLASSO struggled to infer signatures SBS17b and SBS28. Further results presented in Fig 2D, depict that SigProfilerSS and sigLASSO have a higher fitting error of 0.4. Whereas we can observe StarSignDNA, deconstructSigs, MutationalPatterns, and sigfit achieve a low fitting error of 0.1, and stable prediction (S1 Table, S2 Table, and S3 Table).

**Fig 2.**
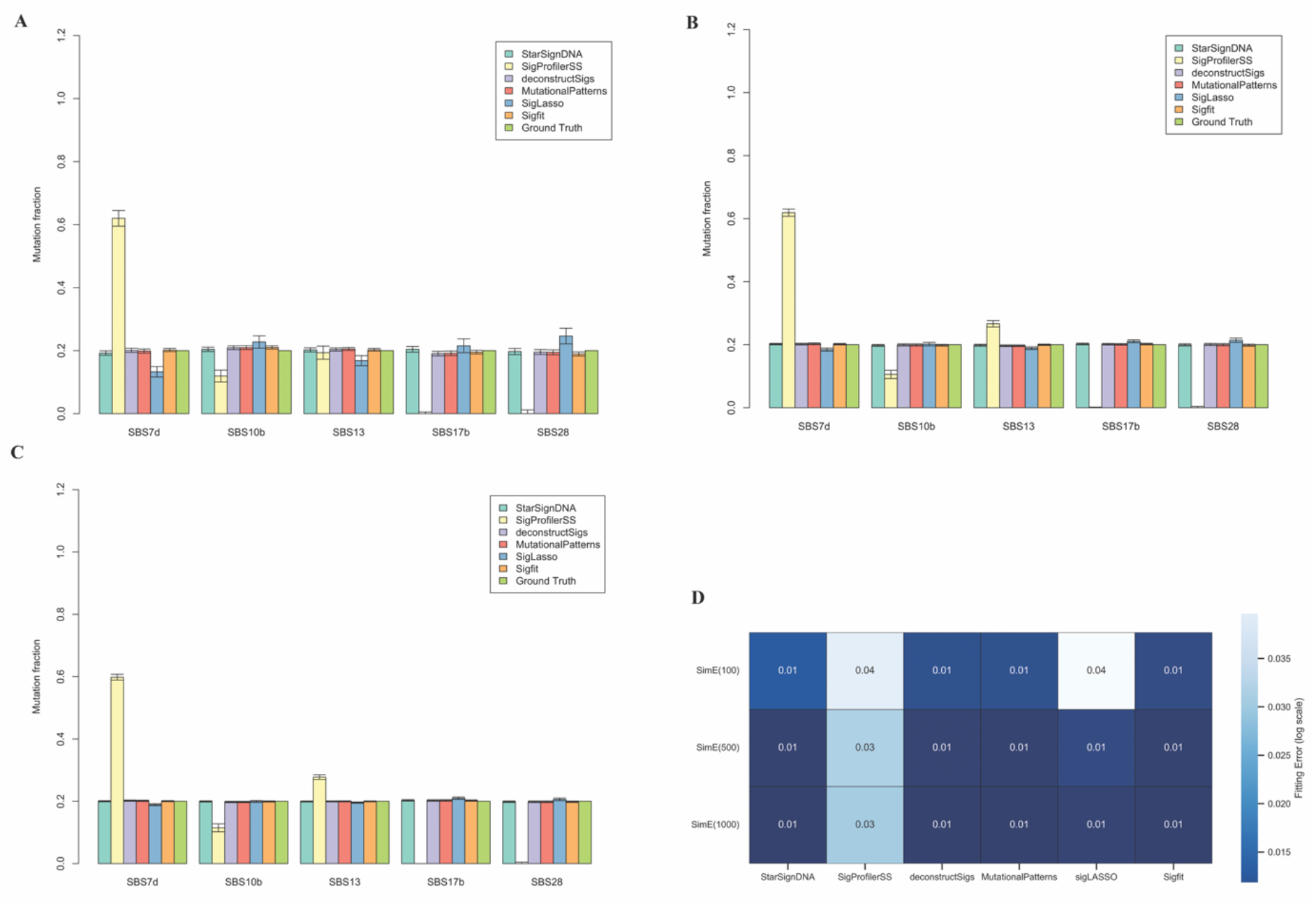
Performance comparison of six mutational signature re-fitting tools. StarSignDNA (light green), SigProfilerSS (light yellow), deconstructSigs (light blue), MutationalPatterns (red), SigLASSO (dark blue), and sigfit (orange) across five different signature types (SBS7d, SBS10b, SBS13, SBS17b, and SBS28) is represented by each set of bars. The vertical lines on top of each bar indicate the variability or uncertainty in the mutation fraction estimation by each tool compared to the known exposure values (Ground Truth) set to 0.25 for each signature. A) shows mutation fraction estimates of 100 samples with 500 mutations per sample. B) shows mutation fraction estimates of 100 samples with 500 mutations per sample. C) shows mutation fraction estimates of 100 samples with 1000 mutations per sample. (D) shows the average fitting error (SimE) across 100 cohorts from each simulation with 100, 500, and 1000 mutations (SimE(100), SimE(500), and SimE(1000)).

Considering that using selected reference signatures (i.e. based on prior knowledge) for re-fitting is commonly recommended ([27, 28]), we evaluated the performance of all the tools with both a limited number of reference signatures and the full collection of signatures. This was done to investigate the impact of the reference signature dimensions on the resulting predictions. The results show that StarSignDNA, MutationalPatterns, deconstructSigs, and sigfit outperform SigProfilerSS and sigLASSO when using a limited number of reference signatures (four in Fig 1 and five in Fig 2). A reliable re-fitting algorithm should ideally maintain good performance regardless of whether a reduced or a larger set of reference signatures is used.

After evaluating the re-fitting algorithm using a limited number of reference signatures, we further evaluated the stability of the six methods with the full COSMIC reference signature v3.4, which contains 86 signatures. Compared to other algorithms, StarSignDNA performed best for simulations with a low number of mutations (Fig 3,S4 Table, S5 Table, and S6 Table). Specifically, it achieved the highest F1 score (0.98), followed by MutationalPatterns (0.96), deconstructSigs (0.95), SigProfilerSS (0.90), sigLASSO (0.77), and sigfit (0.59). For simulations with around 500 mutations, StarSignDNA, MutationalPatterns, and deconstructSigs maintained a high F1 score of 1.00. SigProfilerSS, sigLASSO, and sigfit also performed well with an F1 score of 0.99. Interestingly, all six methods achieved a perfect F1 score (F1 score = 1.00) for simulations with around 1000 mutations. These results suggest that the approach used in StarSignDNA shows particular strength in scenarios where samples have low mutation counts.

**Fig 3.**
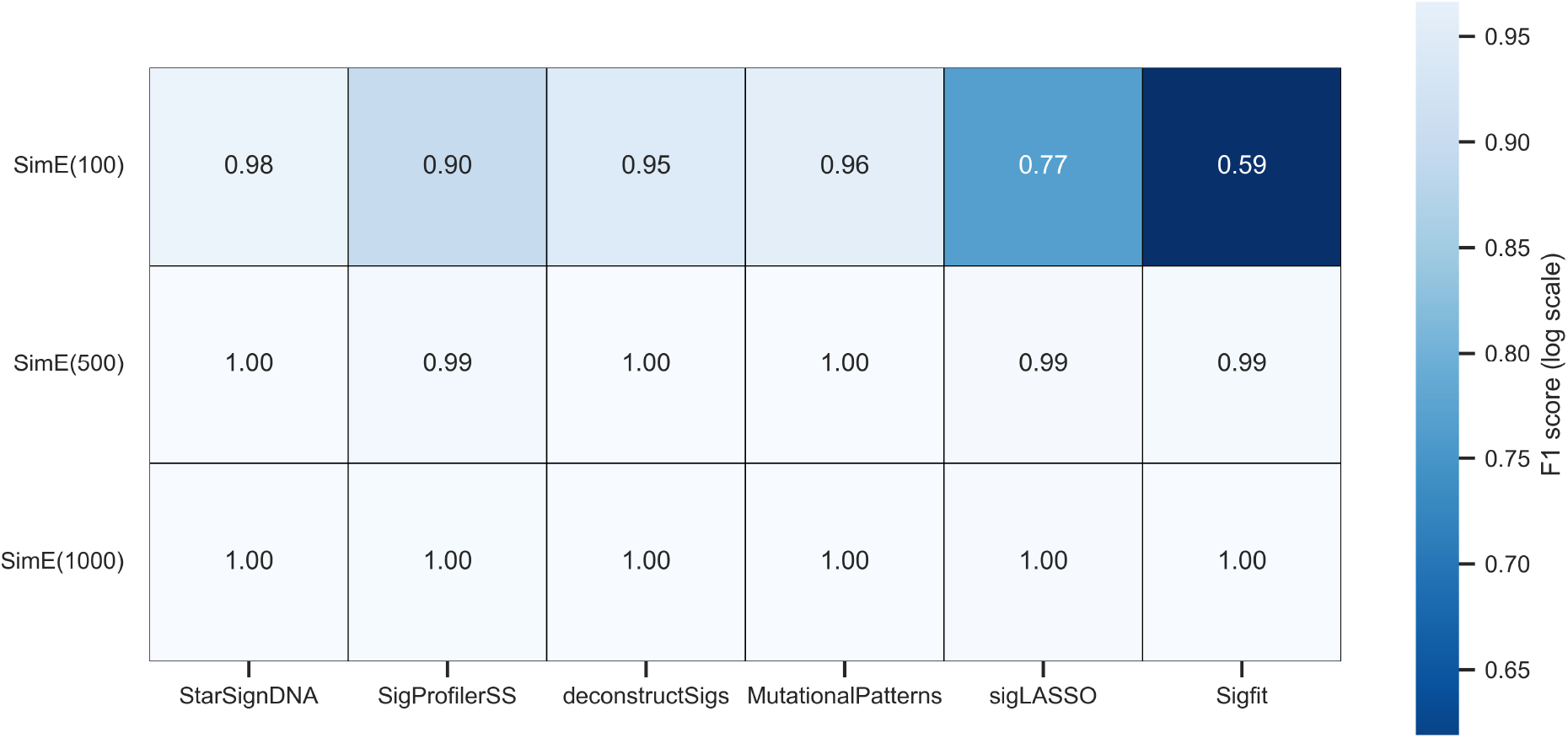
Comparison of F1 scores of six re-fitting methods using SimE(100), SimE(500) and SimE(1000) datasets. The F1 scores were used to evaluate the precision and sensitivity of the signature assignment across the simulations with the threshold exposure of 0.08 (i.e. elements below 0.08 are considered zero).

The COSMIC reference signature collection includes multiple flat signatures, such as SBS3 (related to homologous recombination-based repair) and SBS5 (present in virtually all samples, yet with unknown etiology). These flat signatures exhibit high similarity to other signatures and appear clustered together, making them harder to detect. To evaluate the performance of StarSignDNA on these difficult-to-detect flat signatures, the third category of simulations (Table 3) was used. This category explores the impact of exposure values and mutation counts on the performance of re-fitting algorithms, highlighting the challenges posed by flat signatures.

The analysis began with only five COSMIC v3.4 reference signatures (SBS3, SBS5, SBS40c, SBS92, and SBS94). StarSignDNA, sigfit, MutationalPatterns, and deconstructSigs performed better at inferring flat signatures as compared to SigProfilerSS and sigLASSO, which deviated from the ground truth (Fig 4A-C, S7 Table, S8 Table and S9 Table). The first four methods had the lowest fitting errors, while SigProfilerSS and sigLASSO had the highest fitting errors of 0.1 (Fig 4D). Further analysis using the full set of 86 COSMIC v3.4 signatures on simulations with 100, 500, and 1000 mutations per sample (SimF(100), SimF(500), and SimF(1000)) revealed that for 100 mutations, StarSignDNA achieved the highest F1 score (0.28) followed by SigProfilerSS with a score of 0.26. In contrast, deconstructSigs and sigLASSO had lower scores (0.13). MutationalPatterns and sigfit performed poorly in this scenario, with scores of 0.09 and 0, respectively (Fig 5).

**Fig 4.**
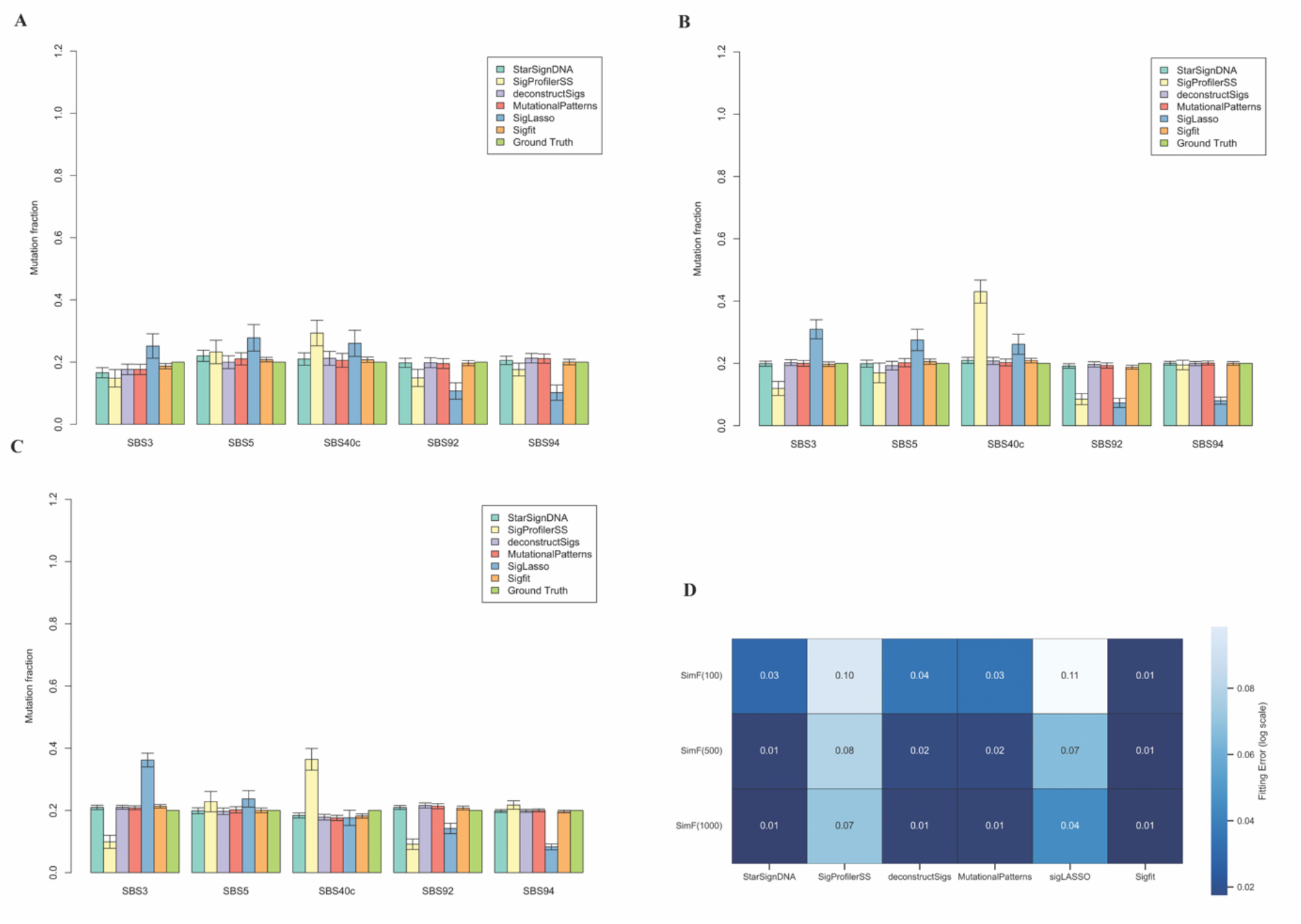
Performance comparison of six mutational signature re-fitting tools. StarSignDNA (light green), SigProfilerSS (light yellow), deconstructSigs (light blue), MutationalPatterns (red), SigLASSO (dark blue), and sigfit (orange) across five different signature types (SBS3, SBS5, SBS40c, SBS92, and SBS94) is represented by each set of bars. The vertical lines on top of each bar indicate the variability or uncertainty in the mutation fraction estimation by each tool compared to the known exposure values (Ground Truth) set to 0.25 for each signature. A) shows mutation fraction estimates of 100 samples with 500 mutations per sample, B) shows mutation fraction estimates of 100 samples with 500 mutations per sample, and C) shows mutation fraction estimates of 100 samples with 1000 mutations per sample. (D) shows the average fitting error (SimF) across 100 cohorts from each simulation with 100, 500, and 1000 mutations (SimF(100), SimF(500), and SimF(1000)).

**Fig 5.**
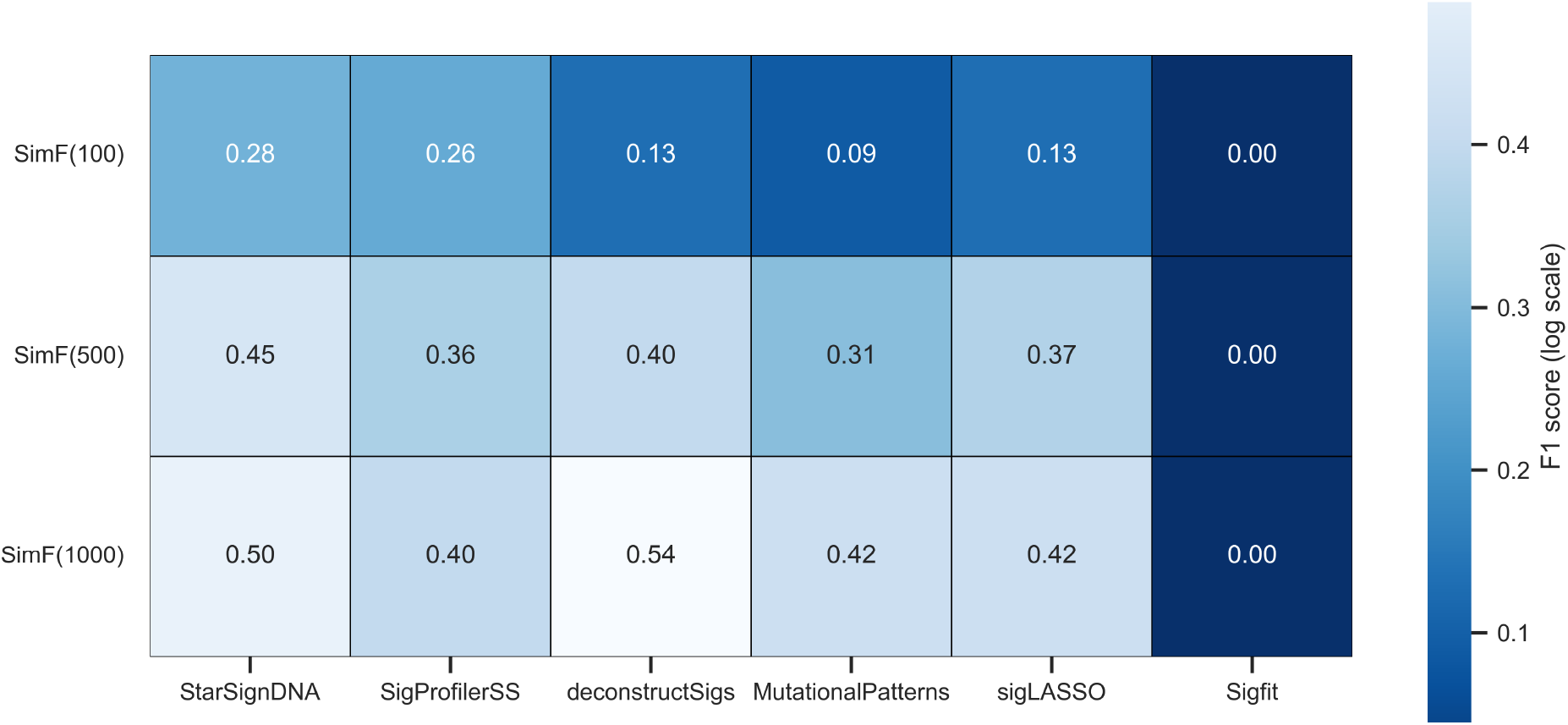
Comparison of F1 scores of six re-fitting methods using SimF(100), SimF(500) and SimF(1000) datasets. The F1 scores were used to evaluate the precision and sensitivity of the signature assignment across the simulations with the threshold exposure of 0.08 (i.e. elements below 0.08 are considered zero).

As the mutation count increased to 500 per sample, StarSignDNA outperformed the other methods with an F1 score of 0.45 followed by deconstructSigs with an F1 score of 0.4. sigLASSO and SigProfilerSS showed improved performance with F1 scores of 0.37 and 0.36 respectively, while MutationalPatterns had a moderate score of 0.31. sigfit performed poorly again, with a score of 0. Moving to 1000 mutations per sample, deconstructSigs performed best with an F1 score of 0.54 followed by StarSignDNA with a score of 0.5, while sigLASSO and MutationalPatterns had scores of 0.42. SigProfilerSS had a score of 0.4, and sigfit continued to score 0. The median Pearson’s correlation coefficient metric was used to compare refit methods. This coefficient measures how well each method fits the observed mutations in individual patients by comparing the observed mutation spectrum to the predicted mutation spectrum for each patient. As shown in Table 4, StarSignDNA exhibited the highest median per-patient correlation across all datasets. SigProfilerSS (Fig 5 and Table 4 (S10 Table, S11 Table, and S12 Table)), indicate that StarSignDNA excels in simulations with lower mutation counts (around 100 and 500), making it suitable for re-fitting analyses of both common and flat signatures. It is important to note that the level of prediction uncertainty is directly related to the number of mutations per sample. In simpler terms, fewer mutations lead to more uncertainty in the sampling process.

We investigated the performance of re-fitting algorithms for flat signatures using downsampled datasets where the mutations are removed randomly from SimF100. As described in (Table 3), SimF50 (50% of mutation removed), SimF70 (30% of mutation removed), SimF80 (20% of mutation removed), SimF90 ((50% of mutation removed)), and SimF100 (Table 3) to assess prediction stability and its correlation with the number of mutations per sample. It aimed to identify the acceptable number of mutations needed for accurate predictions using the full COSMIC v3.4 reference signature set. Results show that StarSignDNA consistently outperformed other methods across these datasets. The highest F1 score for StarSignDNA on SimF50 was 0.22 as shown in (Fig 6). While this is low for high-precision applications, it indicates that around 50 mutations per sample are the threshold for reliable predictions. Below this threshold, predictions become more random and thus, more uncertain. The dimension of the reference signature matrix is crucial for accurate detection with low mutation counts, and prior knowledge of the cancer type to be analyzed guides the selection of relevant signatures S13 Table).

**Fig 6.**
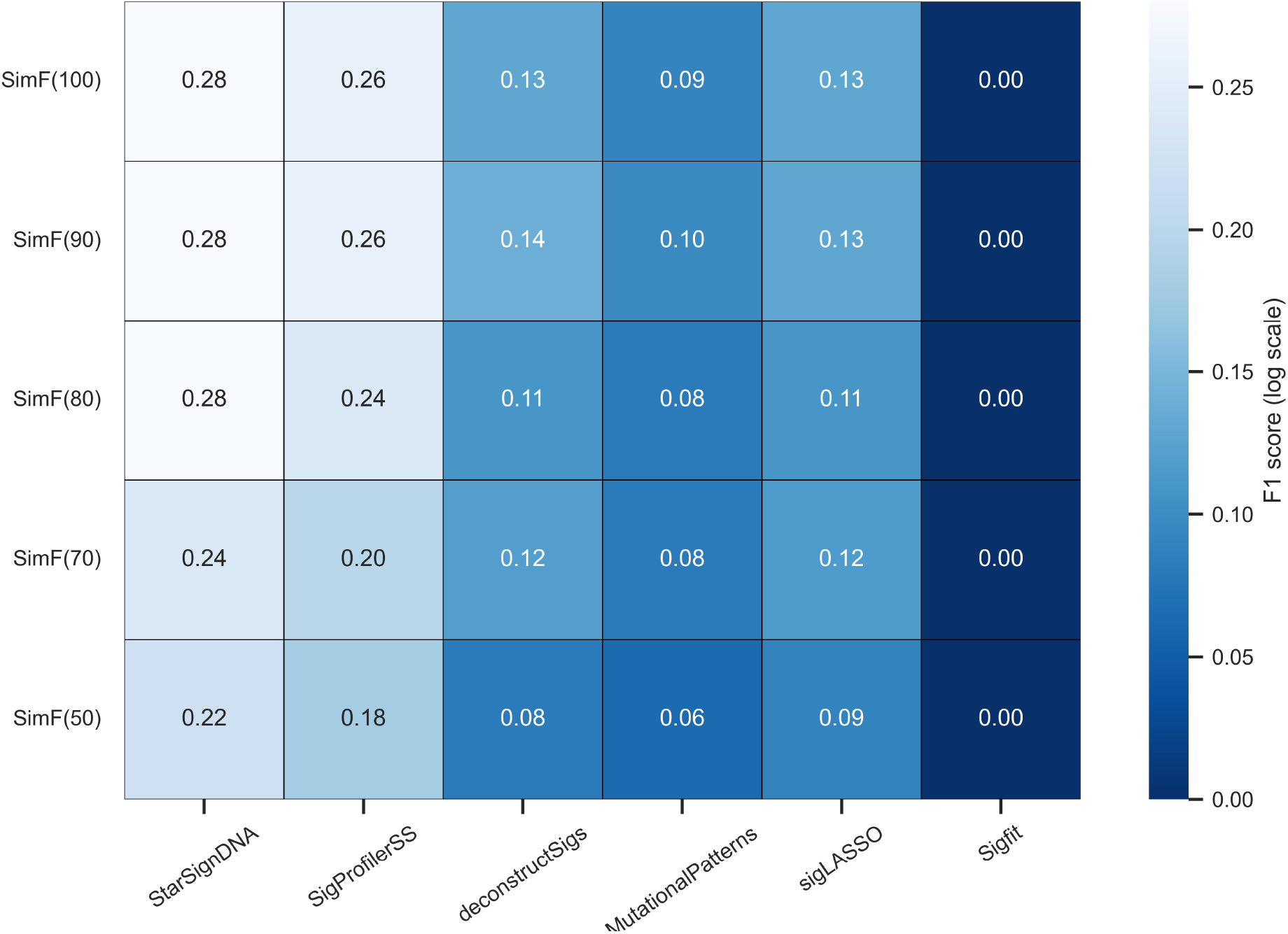
The F1 score was used to evaluate the accuracy of signature assignment across six re-fitting methods. The number of samples for each dataset is 100 with different numbers of mutations. An F1 score threshold of 0.08 was used to evaluate the accuracy. This means elements below 0.08 were considered zero for the signature assignment

The algorithm running time was analyzed as shown in (S5 Figure), and we observed that the total number of reference signatures did not significantly impact computational time. Below 40 reference signatures, all methods except Sigfit demonstrate low running times. When considering over 80 signatures, StarSignDNA, deconstructSigs, MutationalPatterns, and sigLASSO exhibit nearly identical running times.

Lastly, the results from the fourth category of simulations demonstrate that StarSignDNA accurately identified the correct signatures without detecting any additional signatures, using a detection threshold of 6%. In the dataset where SBS17b was simulated, StarSignDNA recovered 78% of SBS17b and 21% of SBS40c. Similarly, in the dataset where SBS92 was simulated, StarSignDNA recovered 71% of SBS92, 13% of SBS40c, and 7% of SBS30 (S6 Figure).

The hyperparameter *λ*, which controls the balance between a good fit to the data and model complexity, is obtained using cross-validation (S7 Figure) from both WES and WGS.

#### StarSignDNA validation on real cohort datasets

We applied StarSignDNA to analyze 8,893 TCGA tumours across 33 cancer types (Fig 7). The result for sigLASSO and deconstructSigs can be found in (S8 Figure and S9 Figure). We used the full COSMIC v3.4 signatures as the reference. The results demonstrate StarSignDNA’s ability to generate sparse signature assignments which can improve prediction owing to lower estimator variance, particularly when dealing with low-count WES datasets.

**Fig 7.**
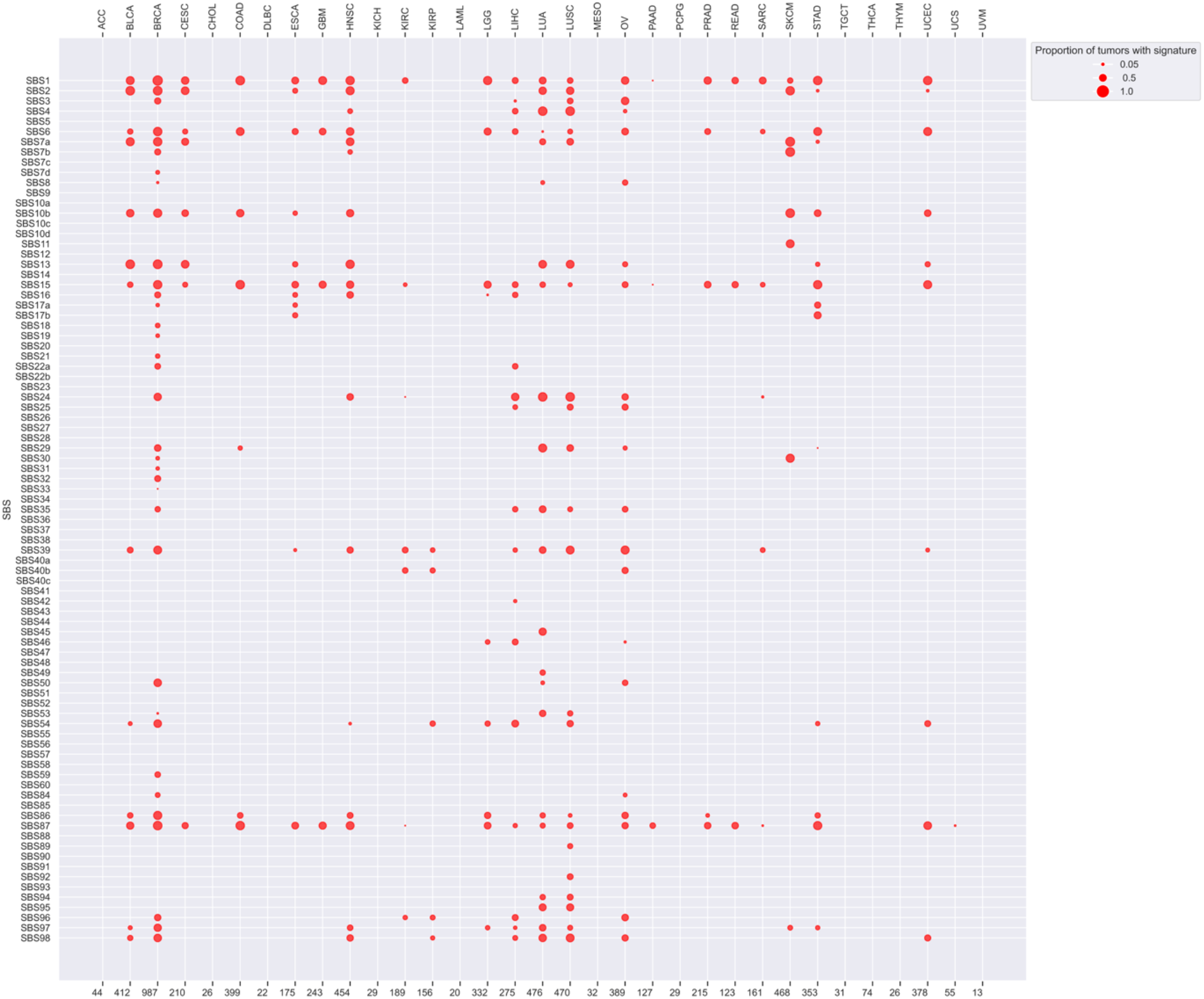
This figure shows the performance of StarSignDNA on 33 TCGA cancer types listed alphabetically along the top x-axis. The size of each dot represents the proportion of signatures within a specific cancer type that exhibits a mutational signature identified by StarSignDNA. The bottom x-axis indicates the number of samples analyzed for each cancer type. A separate panel legend identifies the “active signatures” (those contributing more than 6% of the mutations) in each cancer type using different methods. All 86 COSMIC v3.4 signatures were used as references for the analysis.

#### StarSignDNA validation on single real data samples

In a clinical setting, the detection of mutational signature contributions in single samples is critical. Importantly, the estimation of the uncertainties in the predictions is important for clinical interpretation. StarSignDNA addresses this by incorporating a confidence level for its predictions using bootstrapping. The illustration for a single breast cancer patient sample of TCGA-BRCA is shown in Fig 8.

**Fig 8.**
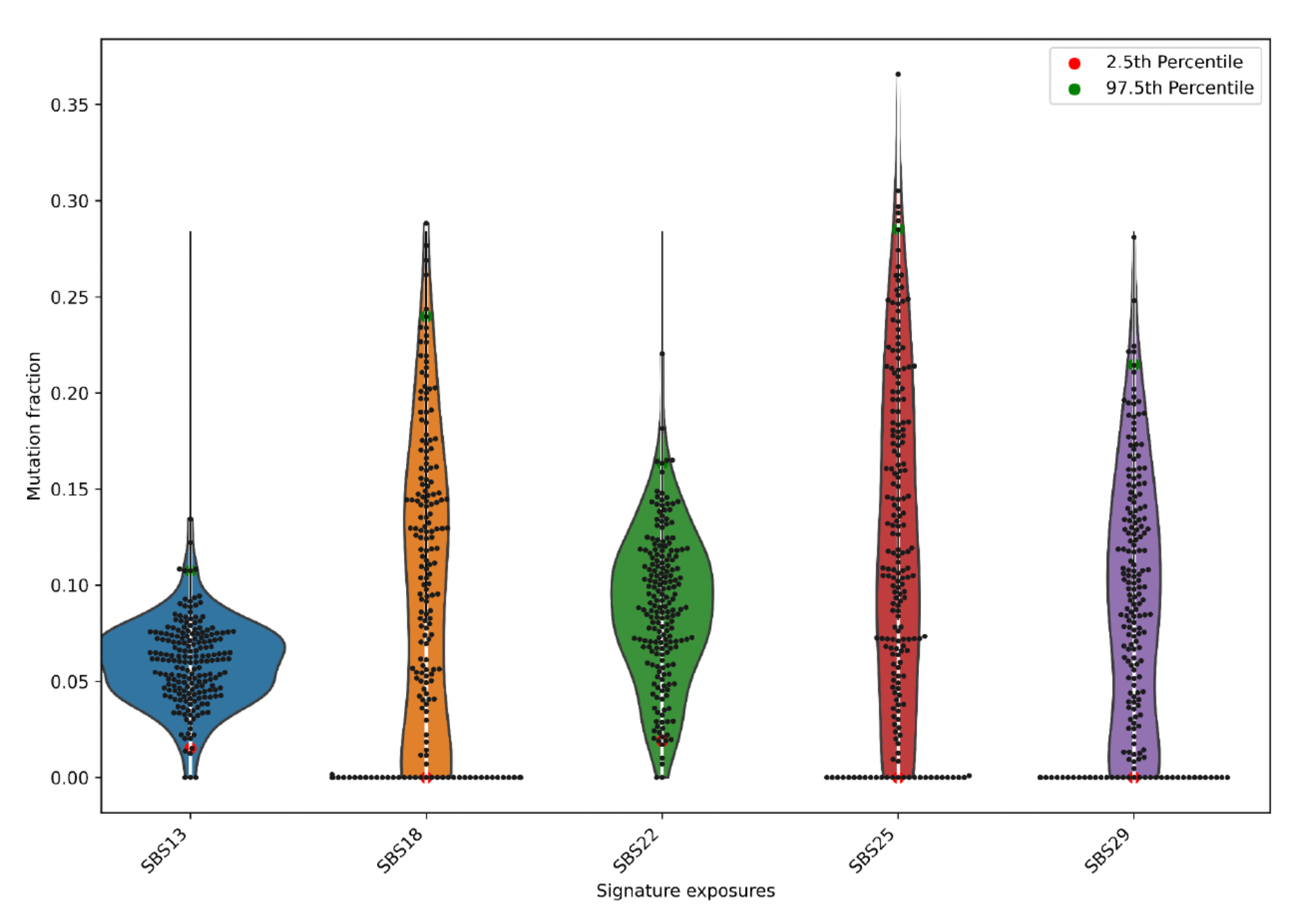
Comparison of mutational re-fitting algorithm for a single sample. The y-axis represents the relative contribution of the exposure, the bar extends from the 2.5% percentile to the 97.5% percentile of the bootstrapped exposure levels obtained from 200 bootstraps, and the x-axis represents the signature exposures.

### *De novo* Mutational signatures

#### StarSignDNA Validation on simulation datasets

StarSignDNA was benchmarked versus other *de novo* algorithms, including SigProfilerExtractor (SigProfiler) [29], an NMF-based method, SparseSignatures [12], a cross-validation method to select both the number of signatures and the shrinkage parameter simultaneously, SUITOR [20], a probabilistic model, and SigneR [8], a Bayesian approach. The benchmarking was performed using the fourth category of simulated data sets. We generated 50 replicate datasets, of 146 patients each, with a set of ground truth mutational signatures being COSMIC SBS1, SBS5, BSB18, and SBS40 (see supporting document). In the *de novo* signature extraction method, identifying the optimal number of signatures (K) is critical. This value of K reflects the underlying biological processes contributing to the mutations. We analyzed a range of values from 2 to 18, allowing each algorithm to select the optimal K value. It is important to note that while the true K for the simulation dataset data is known to be four, additional detectable processes might be present since the simulation dataset data was derived from real-world data. The analysis focused on two key aspects: (i) the number of signatures (K) extracted by each method and (ii) the number of mutational processes each method could recover. Although the cosine similarity threshold can be set to 0.75 when identifying novel signatures [30], in this paper, a stringent threshold of 0.8 was set to define acceptable reconstructed signatures.

Considering the number of extracted signatures (K), all methods identified at least four (K=4), corresponding to the overweighted signatures in the data. SUITOR and SigneR consistently detected only the overweighted signatures across all 50 replicates. SparseSignatures exhibited some variability: K=6 in two replicates, K=4 in 21 replicates, and K=3 in 27 replicates. SigProfiler showed similar behavior with K=4 in 43 replicates, K=3 in three, and K=2 in four replicates. StarSignDNA identified more than four signatures: K=6 in 23 replicates, K=5 in 23 replicates, K=4 in two replicates, and K=3 in two replicates. It’s important to note that finding more signatures reduces residual error and improves data fitting. The primary goal of a *de novo* algorithm goes beyond simply fitting the data. Ideally, it should also identify processes reflecting distinct underlying biological mechanisms. Regarding the residual error, SigneR, SigProfiler, and StarSignDNA achieved the lowest values shown in (Fig 9B). However, (Fig 9A) reveals that SigneR and SigProfiler fail to detect active processes beyond the overweighted ones. In contrast, StarSignDNA and SigneR better reconstruct the original input signatures (Fig 9C). StarSignDNA’s superior accuracy can be attributed to its optimization algorithm and exposure regularization based on learned parameters (*λ*). SparseSignatures, on the other hand, utilize regularization on the signatures themselves, which explains its higher sparsity. Notably, StarSignDNA also exhibits a higher sparsity level compared to other methods (Fig 9D).

**Fig 9.**
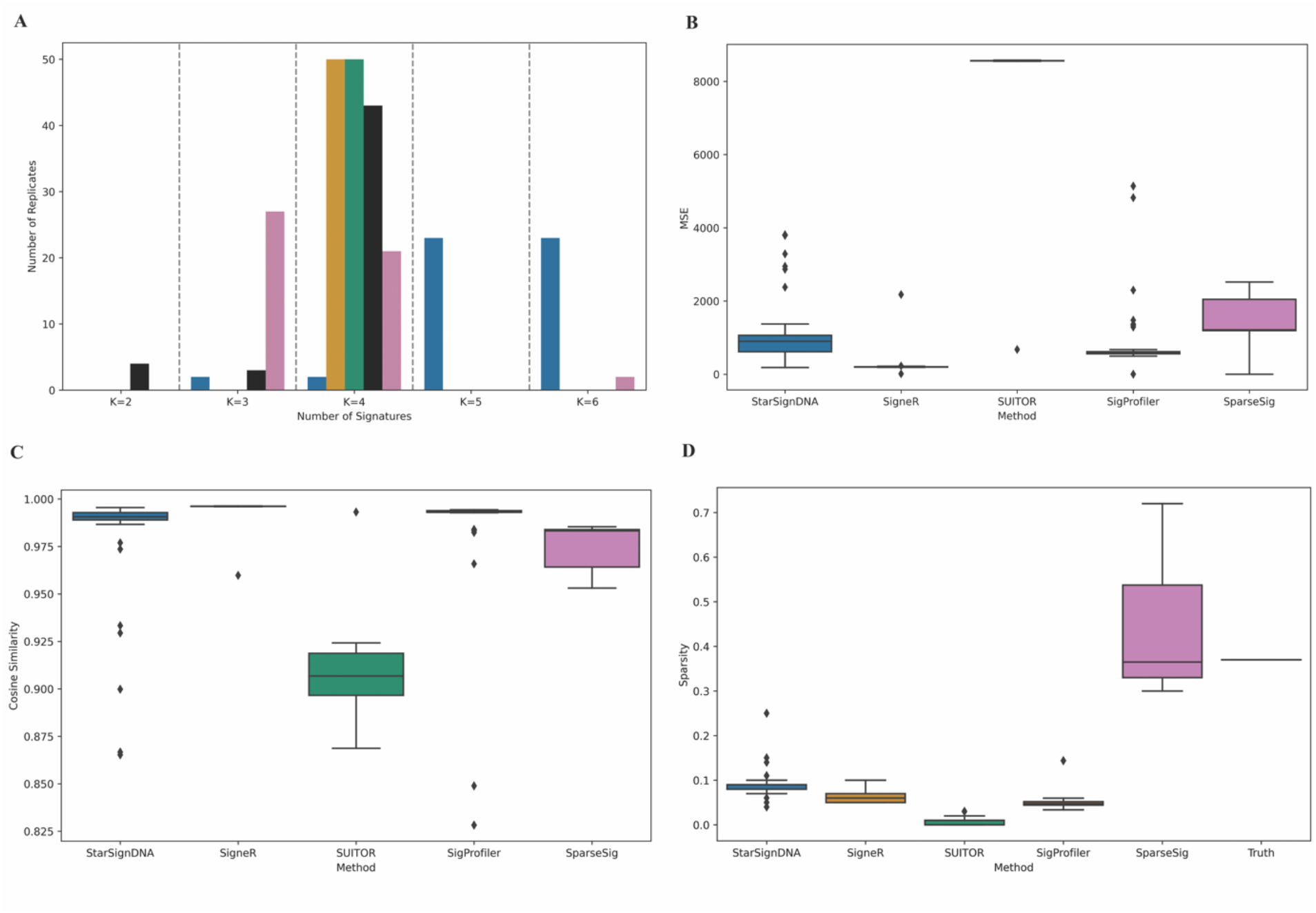
Comparison of StarSignDNA with Other Methods on Simulated Data. A) Number of Signatures Detected (K): This bar chart shows how often each method selected a specific number of signatures (K) across the 50 simulations. The x-axis represents the possible values of K, and the y-axis shows the frequency at which each method chose a particular K. All methods were run on 50 simulated datasets with a ground truth value of K=4 (i.e., the actual number of underlying processes in the data). B) Residual Error: This box plot compares the residual error achieved by each method across the 50 simulations. The residual error is measured using the Mean Squared Error (MSE) between the reconstructed and original count matrices. Lower MSE indicates better reconstruction. C) Cosine Similarity: This box plot shows the cosine similarity between the reconstructed and original input signatures for each method across the 50 simulations. A higher cosine similarity value signifies a more accurate reconstruction of the original signatures. D) Sparsity of Signatures: This box plot compares the sparsity of the signatures generated by each method across the simulations. Sparsity is measured as the proportion of elements in the signature matrix with values below a threshold of 10^3^. Higher sparsity indicates a higher number of zeros within the signatures. The data underlying these analyses can be found in (S6 Table).

StarSignDNA detected SBS1 in 38 replicates, SBS5 in 40 replicates, SBS18 in 31 replicates, SBS40 in 48 replicates, and SBS6 in two replicates. SigneR detects SBS1 in 49 replicates, SBS5 in 49 replicates, and SBS18 in 37 and 49 replicates. SUITOR detects SBS1 in 40 replicates, SBS5 in 49 replicates, SBS18 in 35 replicates, and SBS40 in 49 replicates. SparseSignatures detect SBS1 in 22 replicates, SBS40 in 49 replicates, and SBS60 in nine replicates. Sigprofiler detected SBS1 in 49 replicates, SBS3 in three replicates, SBS5 in 42 replicates, SBS18 in 49 replicates, SBS40 in 46 replicates, and SBS16 in seven replicates. As shown in (Fig 10), it indicates that StarSignDNA outperforms the other algorithms by capturing signature SBS3 in 36 replicates compared to other methods (SigProfiler: three replicates, SUITOR: one replicate, SigneR, and SparseSignatures: 0 replicate). This is likely because SBS3 is a difficult-to-detect signature due to its high similarity to other signatures. This finding aligns with (Fig 9A), which demonstrated StarSignDNA’s ability to detect less frequent and potentially more challenging processes. It’s further corroborated by the re-fitting results in (Fig 5-6). The superior performance of StarSignDNA in detecting SBS3 might be attributed to its grid-search algorithm. This algorithm employs a combination of cross-validation and the probability mass function to effectively select the optimal number of active processes within the data.

**Fig 10.**
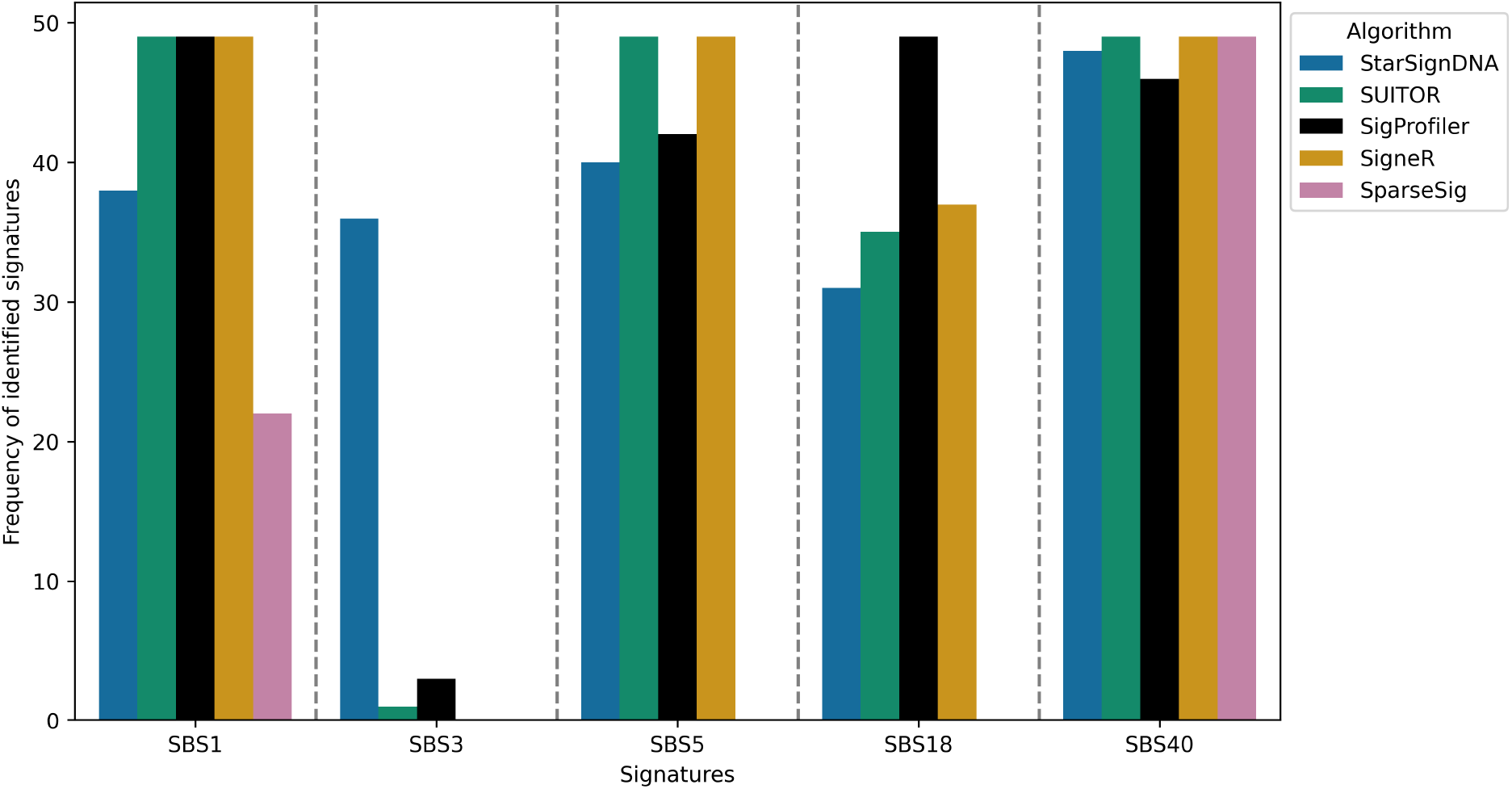
*In silico* evaluation of StarSignDNA and other methods. The number of replicates in which a given signature (of a total of 4 signatures) will be detected by each method.

#### De novo mutational signatures validation on real datasets

*De novo* mutational signature extraction was validated on real data from whole-genome sequencing (WGS) of tumours across five cancer types (Table 5). These cancer types include breast, skin, prostate, kidney, and pancreatic. The data consisted of 1,049 tumours from PCAWG and 2,924 tumours from HMF.

#### Mutational signature on primary tumour data (PCAWG)

All five algorithms were run on the same datasets, allowing them to search for the optimal number of signatures (K) in the interval from 2 to 18 as illustrated for skin cancer (Table 6). In this paper, a *higher signature* refers to a detected *de novo* signature with a cosine similarity equal to or greater than 0.8. SigneR detects 18 Signatures of which only eight identified to be higher signatures and the rest are considered as noise. Similarly, the package SparseSignatures detects 11 signatures out of which only one is identified to be a higher signature while the rest was noise. StarSignDNA and SUITOR detected four signatures out of which three were identified as higher signatures and both the packages had a better prediction ratio of 0.75. This suggests that StarSignDNA and SUITOR identified more relevant signatures with less noise compared to SigneR and SparseSignatures, even though they used a lower number of signatures (K).

**Table 6.** Comparison of signatures predicted by five *de novo* methods on PCAWG skin tumour. Sparsity is measured as the fraction of cells in the matrix with a value = 0. Cosine similarity is measured as the mean of the reconstructed matrix and the original count matrix. Median per-patient correlation is measured as the median Pearson’s correlation coefficient between the observed mutation section and the predicted mutational spectrum for each patient, indicating how well each method fits the observed mutations in individual patients. The prediction ratio is calculated between the total number of signatures, as detected with cosine similarity above 0.8, and the total number of signatures detected by each method. (The table header stands for; CS: Cosine Similarity; NR: Number signatures (K=2 … 18), SigCos: Number of higher signature; Median: Median per-patient correlation

| Method | CS | NR | MSE | SigCos | Pred Ratio | Sparsity | Median |
| --- | --- | --- | --- | --- | --- | --- | --- |
| StarSignDNA | 0.99 | 4 | 74897 | 3 | 0.75 | 0,146 | 0.993 |
| SigneR | 0.99 | 18 | 6067 | 8 | 0.44 | 0.000 | 0.99 |
| SigProfiler | 0.99 | 5 | 26215 | 3 | 0.60 | 0.000 | 0.986 |
| SUITOR | 0.72 | 4 | 400693 | 3 | 0.75 | 0.000 | 0.719 |
| SparseSignatures | 0.99 | 11 | 82289 | 1 | 0.09 | 0,160 | 0.992 |

Unlike simulated data with known true signatures, real-world data lacks this information. However, we can evaluate if the detected signatures correlate highly with the observed mutations. As presented in Table 6, the median per-patient correlation coefficient (measured by Pearson’s correlation) was used to assess how well each method fits the observed mutations in individual patients. This coefficient compares the observed mutation spectrum to the predicted one, for each patient. StarSignDNA demonstrated the best fit to individual patient mutation counts (median correlation = 0.993), followed by SparseSignatures (0.992). This indicates that StarSignDNA and SparseSignatures captured the underlying mutational processes more effectively in these real-world cancer genomes.

Additionally, *de novo* signatures detected by five algorithms are presented in Fig 11. The corresponding cosine similarity to COSMIC reference signatures is available in supplementary tables for breast (S14 Table and S24 Table), skin (S15 Table and S25 Table), pancreatic (S16 Table and S26 Table), prostate (S17 Table and S27 Table), and kidney (S18 Table and S28 Table).

**Fig 11.**
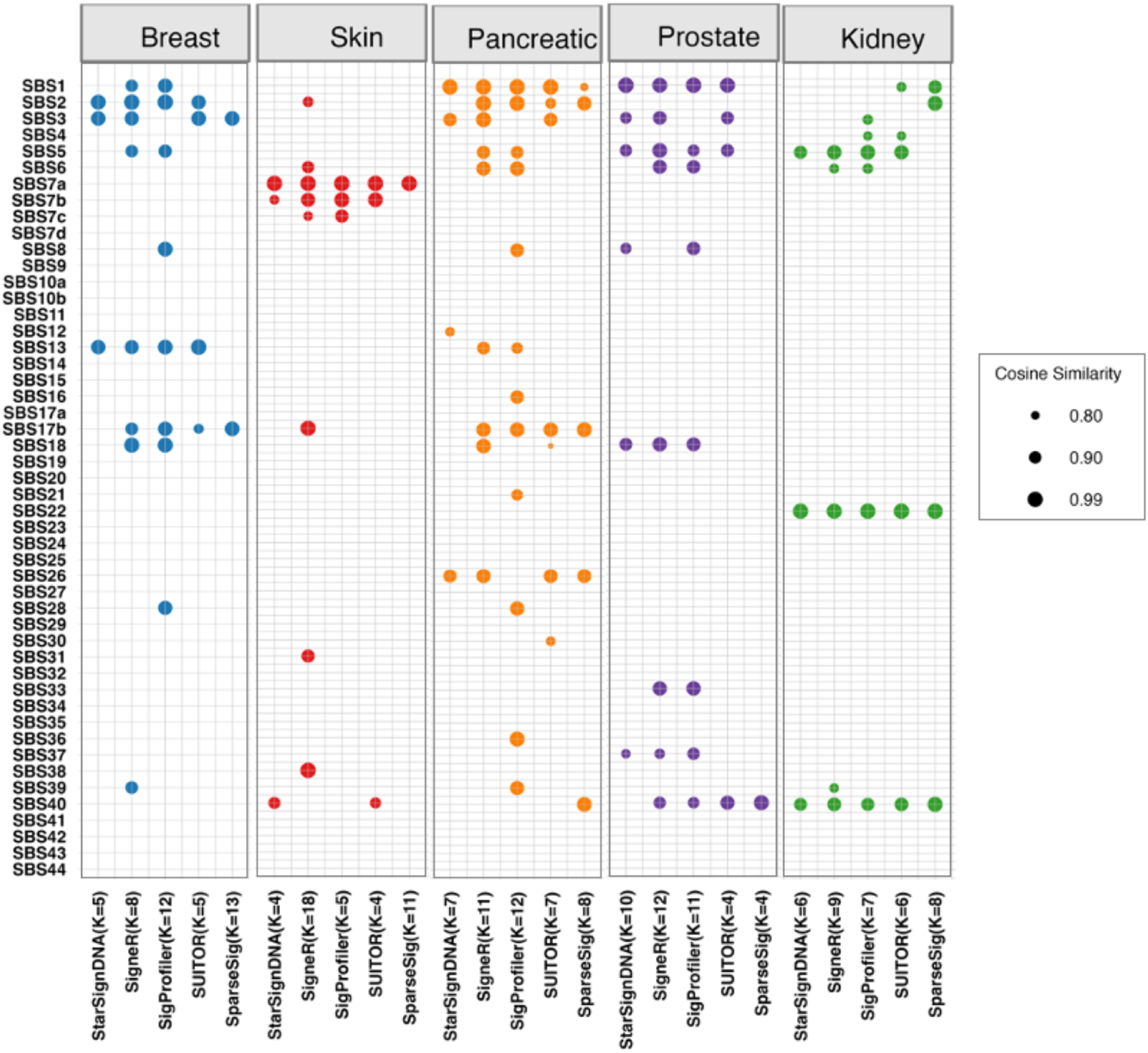
Mutational signature results of five cancer types in PCAWG. The cosine similarities are between *de novo* signatures and COSMIC signatures. Only the matched pairs are shown (with cosine similarity *>* 0.8). The higher the cosine similarity, the better the match to the COSMIC signature profile. The cosine similarity equivalent to one denotes a perfect match. The value of K represents the number of *de novo* signatures discovered by each method.

Delving into the concept of sparsity which refers to the number of inactive elements (zeros) within a matrix, we observed that StarSignDNA achieves a lower value (0.15) compared to SparseSignatures (0.16). In the context of signatures, a lower sparsity indicates that more mutational signatures are present potentially contributing to the observed mutations. Interestingly, both packages (StarSignDNA and SparseSignatures) employ regularization techniques to prevent overfitting, but they achieve this in distinct ways. SparseSignatures directly enforces sparsity by driving elements in the signature matrix towards zero, potentially eliminating weak but relevant signatures. In contrast, StarSignDNA focuses on the exposure values, which define the contribution of each signature. Shrinking these exposure values indirectly achieves sparsity while potentially retaining weak signatures within the exposure values themselves. This difference in regularization strategies might explain why StarSignDNA captures a wider range of mutational processes, leading to potentially more interpretable biological insights.

Overall, these results suggest that StarSignDNA performs well on primary WGS tumour data. It achieves high-fitting performance, identifies relevant signatures with less noise, and extracts sparser signatures more likely to represent distinct mutational processes.

#### Detection mutational signatures on metastatic (HMF)

The same approach used for the primary tumour data was applied to the metastatic data, to assess the prediction performance of five methods, using a dataset of 2,924 metastatic samples. As shown in Table 7 for skin cancer, SparseSignatures fit the counts of individual patients better (0.995), followed by StarSignDNA (0.994). The signatures predicted by SigneR (K=17) and SigProfiler (K=16) appear to contain more noise. StarSignDNA (K=3) and SUITOR (K=3) show a better prediction ratio of 1.00 followed by SparseSignatures with a prediction ratio of 0.67.

**Table 7.** Comparison of signatures predicted by five *de novo* methods on HMF skin tumour. Sparsity is measured as the fraction of cells in the matrix with a value = 0. Cosine similarity is measured as the mean of the reconstructed matrix and the original count matrix. Median per-patient correlation is measured as the median Pearson’s correlation coefficient between the observed mutation section and the predicted mutational spectrum for each patient, indicating how well each method fits the observed mutations in individual patients. The prediction ratio is calculated between the total number of signatures, as detected with cosine similarity above 0.8, and the total number of signatures detected by each method. (The table header stands for; CS: Cosine Similarity; NR: Number signatures (K=2 … 18), SigCos: Number of higher signature; Median: Median per-patient correlation)

| Method | CS | NR | MSE | SigCos | Ratio | Sparsity | Median |
| --- | --- | --- | --- | --- | --- | --- | --- |
| StarSignDNA | 1.00 | 3 | 97837 | 3 | 1.00 | 0.041 | 0.994 |
| SigneR | 1.00 | 17 | 7713 | 8 | 0.47 | 0.000 | 0.99 |
| SigProfiler | 1.00 | 12 | 73239 | 8 | 0.67 | 0.018 | 0.98 |
| SUITOR | 0.88 | 3 | 606961 | 3 | 1.00 | 0.000 | 0.98 |
| SparseSignatures | 0.99 | 3 | 150344 | 2 | 0.67 | 0.131 | 0.995 |

Analyzing actual signatures extracted when considering the higher signatures, SigneR detected 17 signatures and only eight higher signatures. SigProfiler detected 12 signatures and only eight higher signatures. SparseSignatures detected three signatures, and two higher signatures. SUITOR detected three signatures and three higher signatures. StarSignDNA detected three signatures and three higher signatures.

Furthermore, *de novo* signatures detected by the five algorithms are presented in Fig 12. The corresponding cosine similarity to COSMIC reference signatures is available in supporting tables for breast (S19 Table and S29 Table), skin (S20 Table and S30 Table), pancreatic (S21 Table and S31 Table), prostate (S22 Table and S32 Table), and kidney (S23 Table and S33 Table) Like the primary tumour analysis findings, StarSignDNA and SparseSignatures achieved a high median per-patient correlation, indicating good fits to the observed mutations. Additionally, StarSignDNA and SUITOR maintained a higher prediction ratio, suggesting they identified more relevant signatures with less noise.

**Fig 12.**
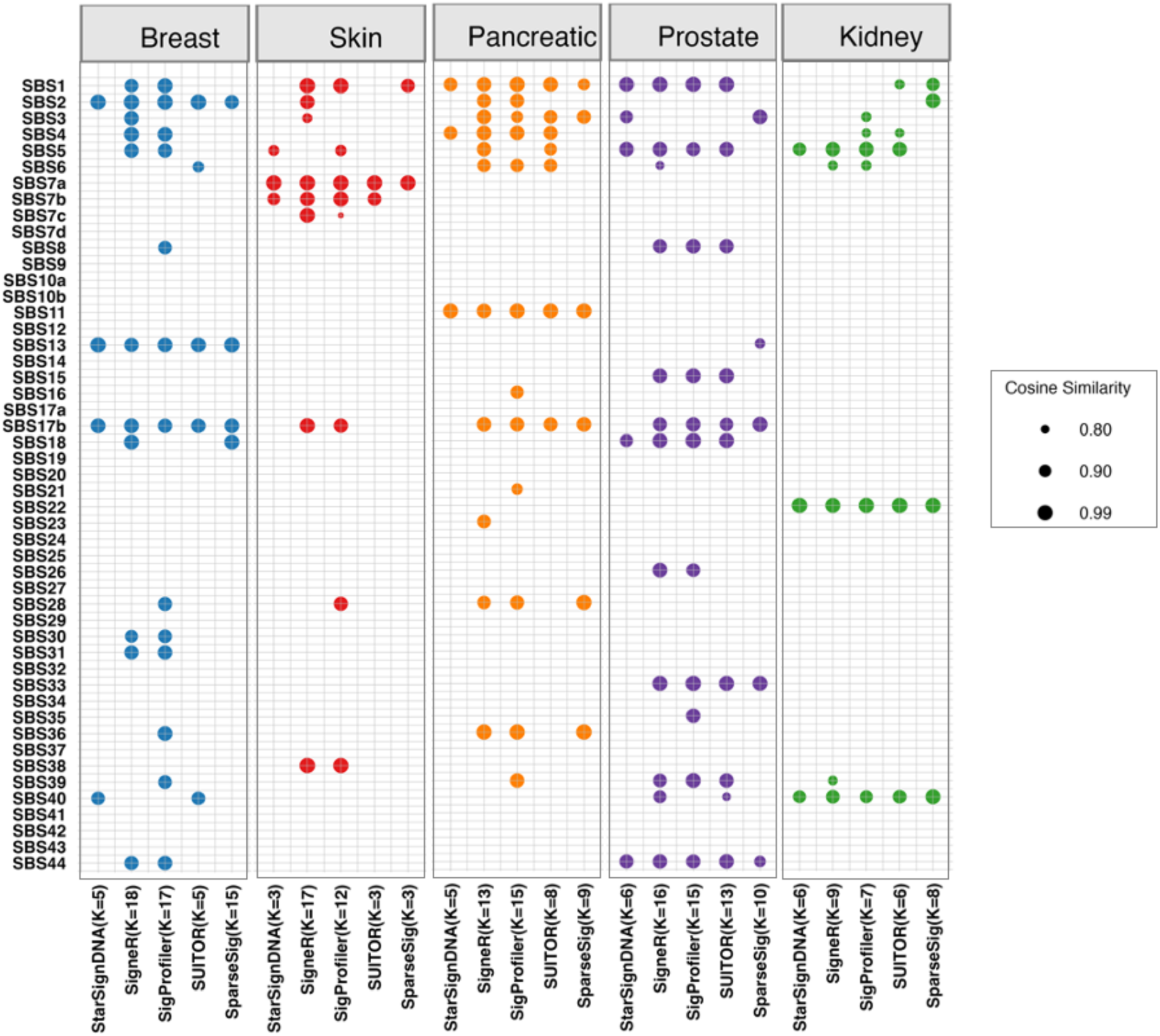
Mutational signature results of five metastatic cancer types from HMF data. The cosine similarities between *de novo* signatures and COSMIC signatures are shown. Only the matched pairs are shown (with cosine similarity *>* 0.8). The higher the cosine similarity, the better the match to a COSMIC signature profile. A cosine similarity equivalent to one denotes a perfect match. The value of K represents the number of *de novo* signatures discovered by each method.

The result on primary tumour and metastatic data shows prediction stability StarSignDNA methods.

### Evaluation of Mutational Signature Quality Using COSMIC Reference

To assess the quality of signatures identified by the five algorithms, this analysis focused on breast and skin cancer types using the COSMIC v3.4 database as the reference. The results of each cancer type and the algorithm included a signature matrix and exposure matrix. The evaluation involved computing the similarity (*Q*_*ij*_) between each pair of signature matrices, matching K signatures from one algorithm with another, and calculating a quality score (q) based on cosine similarity. *Q*_*ij*_ was the maximum quality score across all pairwise comparisons for a specific cancer type. Higher *Q*_*ij*_ values indicate better alignment. StarSignDNA (Fig 13A and Fig 13B) achieved the highest signature quality for both breast (0.55) and skin (0.37) cancer, indicating greater similarity to the COSMIC reference signatures.

**Fig 13.**
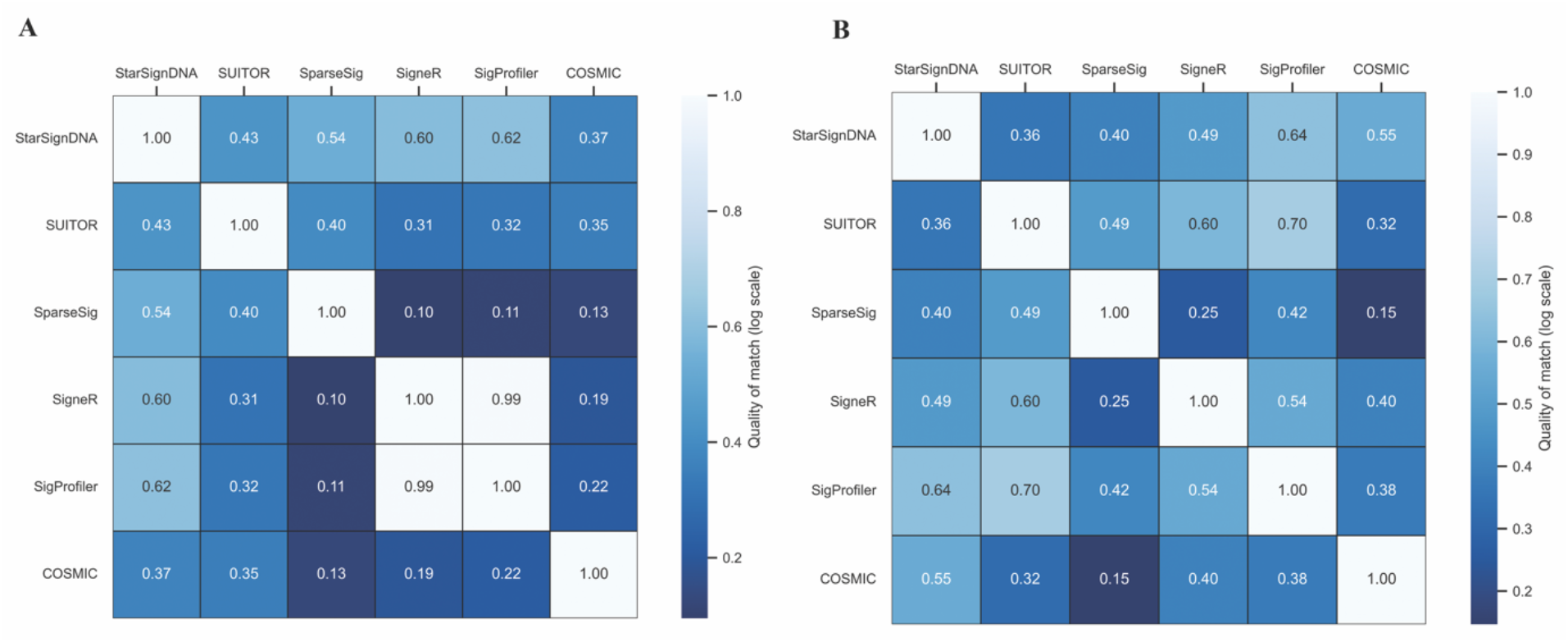
Comparison of *de novo* Signature Quality by Algorithm. A) PCAWG Skin Cancer compares the quality of *de novo* signatures identified by different algorithms for PCAWG skin cancer data. The analysis considers a set of 11 mutational signatures: SBS1, SBS2, SBS5, SBS7a, SBS7b,SBS7c, SBS7d, SBS13, SBS17a, SBS17b, and SBS40. B) PCAWG Breast Cancer compares the quality of *de novo* signatures identified by different algorithms for PCAWG breast cancer data. The analysis considers a set of 12 mutational signatures: SBS1, SBS2, SBS3, SBS5, SBS8, SBS13, SBS17a, SBS17b,SBS18, SBS37, SBS40, and SBS41.

## Discussion

Over a decade after the Alexandrov et al., [17] publication of mutational signatures, various algorithms have emerged, producing similar but not identical results, raising questions about prediction accuracy and biological validity. A key challenge is applying insights from large-scale signature studies to datasets with few mutations and samples. Despite the suitability of decomposition methods for both large and small datasets, challenges remain. These challenges include preventing overfitting and underfitting in low mutation samples, achieving optimal variable selection and shrinkage with full COSMIC reference signatures, and ensuring the biological interpretability of predictions. This paper introduces StarSignDNA, a new algorithm for identifying mutational signatures from cancer sequencing data. StarSignDNA utilizes the L1-penalized estimation algorithm developed by Goeman [31], which employs directional Taylor approximations to maximize the penalized likelihood function.

Since StarSignDNA incorporates L1-regularisation, it achieves more interpretable and accurate identification of mutational signatures from cancer sequencing data.

To evaluate StarSignDNA’s effectiveness, its performance was compared against six established re-fitting algorithms and five *de novo* signature extraction algorithms using both simulation datasets (2,046 samples) and real cancer data (12,201 samples).

StarSignDNA excels in scenarios with low mutation counts, balancing the number of signatures detected and the discovery of true mutational signatures.

The enhanced performance of StarSignDNA can improve our understanding of the key drivers of cancer initiation and the factors affecting the detection of mutational signatures and latent mutational processes.

StarSignDNA demonstrates significantly better performance in re-fitting scenarios, especially with low mutation counts. By applying L1-regularization, StarSignDNA reduces prediction variance and induces sparsity, resulting in more precise predictions than L2-regularization. This approach leads to a better fit of observed mutations in individual patients, and lower fitting errors (Fig 1 to Fig 7). Analysis of four downsampled datasets (Fig 6) shows that the mutation count cutoff is proportional to the dimension of the COSMIC reference matrix used for fitting. Table 4 further underscores the improved fit of observed mutations in patients, offering substantial value for clinicians and researchers. Additionally, StarSignDNA provides confidence level estimates for single sample predictions, helping decision-makers understand the range of likely true parameter values to make informed decisions about mutational processes, even in scenarios with inherent uncertainty. It also demonstrates improved detection of challenge signatures and a better prediction ratio. One area for improvement in the *de novo* method is reducing the computational runtime of the grid search for selecting optimal values for *λ* and *K*.

StarSignDNA is a powerful tool for mutational signature detection, excelling in both re-fitting and *de novo* analyses. Its re-fitting module addresses mutation sampling uncertainty. For *de novo* analysis, StarSignDNA uses the probability mass function as the cost function, achieving better alignment between detected signatures and those with high similarity (cosine similarity *>* 0.8) to known signatures.

In summary, we demonstrated that StarSignDNA is a reliable tool for both *de novo* and re-fitting mutational signature analysis. It offers unique features, such as prediction confidence and customizable reference signatures for the re-fitting analysis while for the *de novo* it gives the prediction ratio. These enhancements improve upon current methods and provide valuable insights for genomics and oncology research.

## Supporting information

**S1 Figure.**
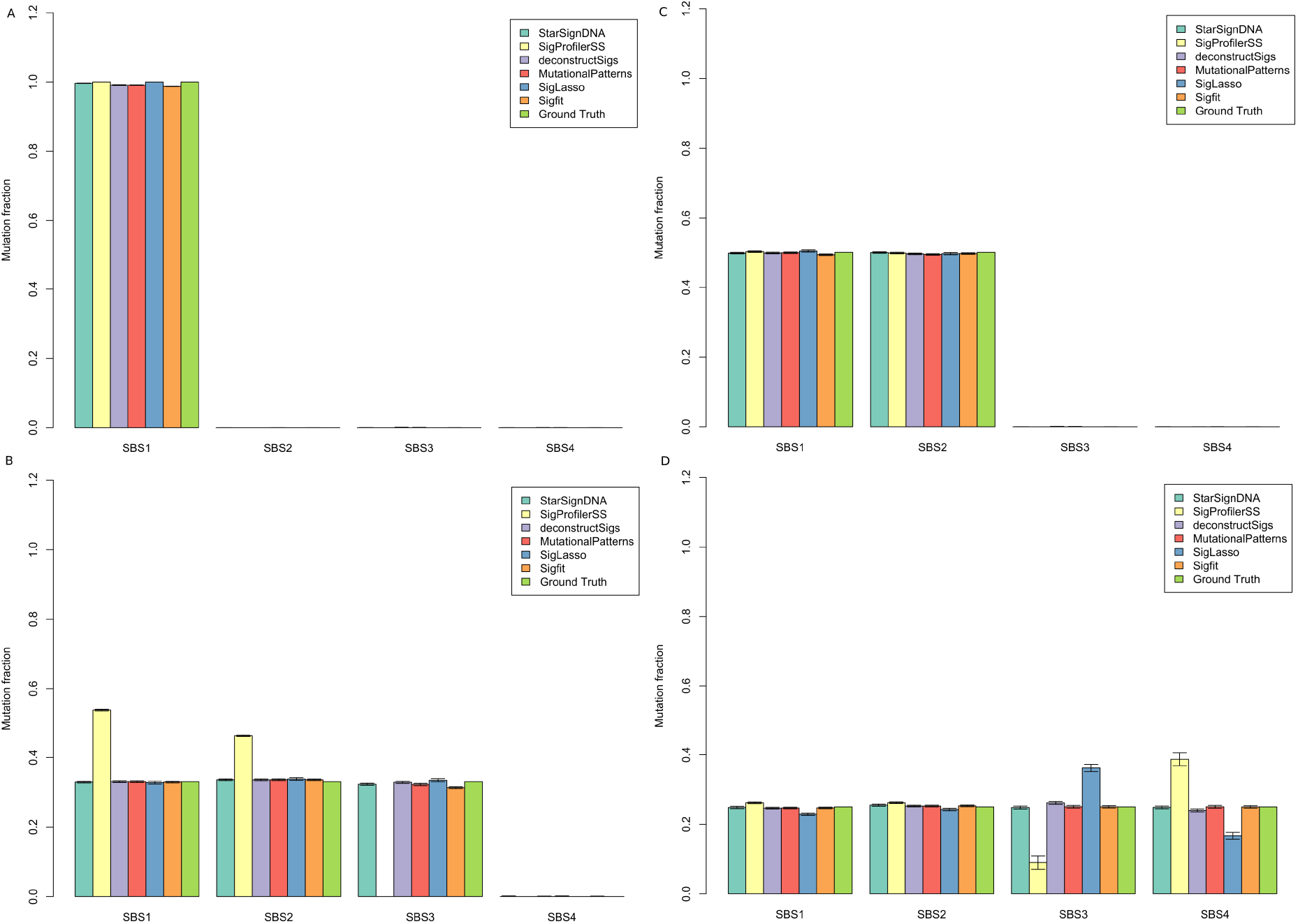
Performance comparison of six mutational signature re-fitting tools. StarSignDNA (light green), SigProfilerSS (light yellow), deconstructSigs (light blue), MutationalPatterns (red), SigLASSO (dark blue), and sigfit (orange) across four different signature types (SBS1, SBS2, SBS3, and SBS4) is represented by each set of bars. The vertical lines on top of each bar represent the average exposure of 100 simulated samples (approximately 500 mutations each) indicating the variability or uncertainty in the mutation fraction estimation by each tool compared to the known exposure values (Ground Truth). A) Only SBS1 has a true exposure of 1, while SBS2, SBS3, and SBS4 have a true exposure of 0. B) SBS1 and SBS2 have true exposures of 0.5, while SBS3 and SBS4 have true exposures of 0. C) SBS1, SBS2, and SBS3 each have true exposures of 0.333, while SBS4 has a true exposure of 0. D) All four signatures have an equal true exposure of 0.25.

**S2 Figure.**
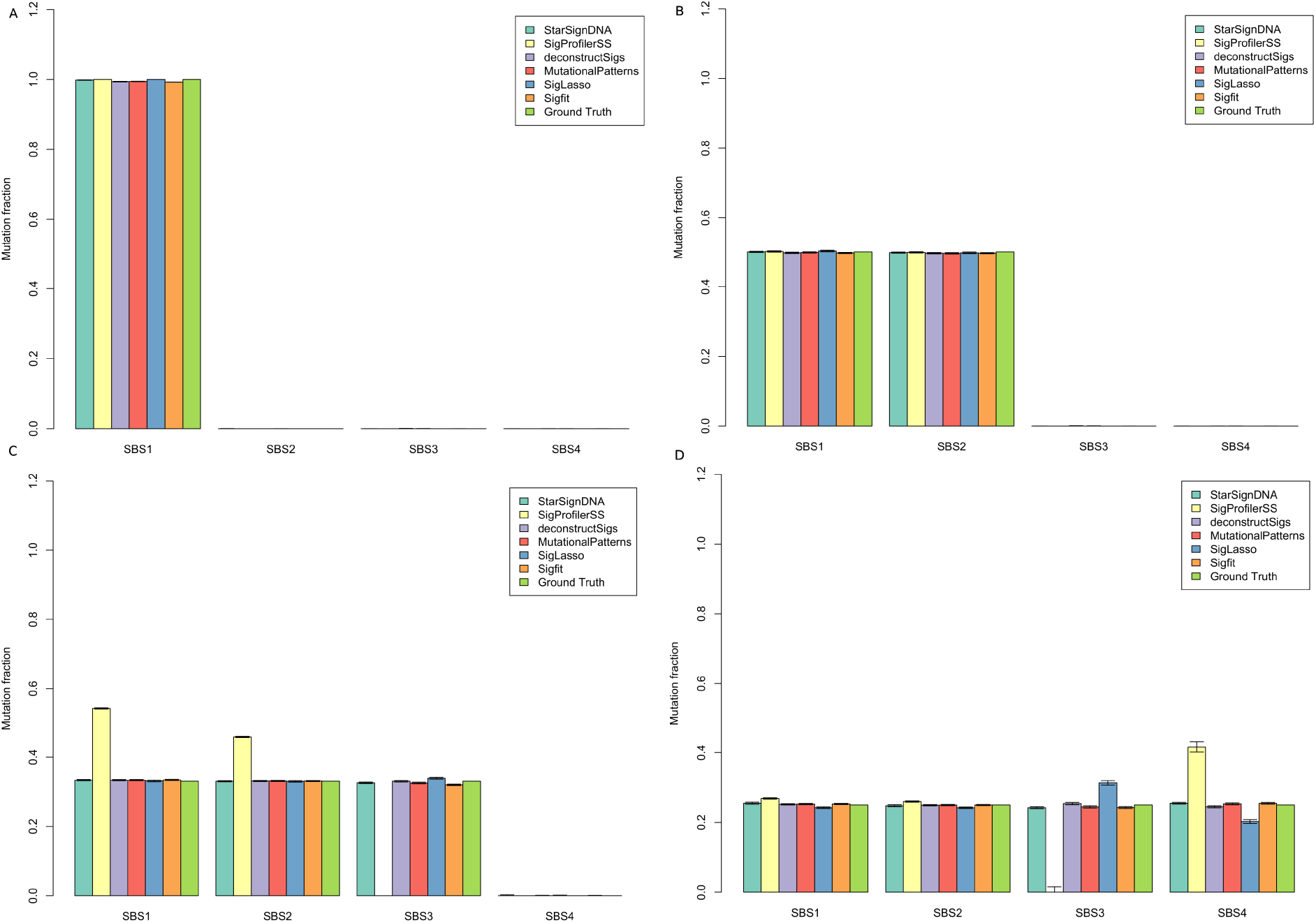
Performance comparison of six mutational signature re-fitting tools. StarSignDNA (light green), SigProfilerSS (light yellow), deconstructSigs (light blue), MutationalPatterns (red), SigLASSO (dark blue), and sigfit (orange) across four different signature types (SBS1, SBS2, SBS3, and SBS4) is represented by each set of bars. The vertical lines on top of each bar represent the average exposure of 100 simulated samples (approximately 1000 mutations each) indicating the variability or uncertainty in the mutation fraction estimation by each tool compared to the known exposure values (Ground Truth). A) Only SBS1 has a true exposure of 1, while SBS2, SBS3, and SBS4 have a true exposure of 0. B) SBS1 and SBS2 have true exposures of 0.5, while SBS3 and SBS4 have true exposures of 0. C) SBS1, SBS2, and SBS3 each have true exposures of 0.333, while SBS4 has a true exposure of 0. D) All four signatures have an equal true exposure of 0.25.

**S3 Figure.**
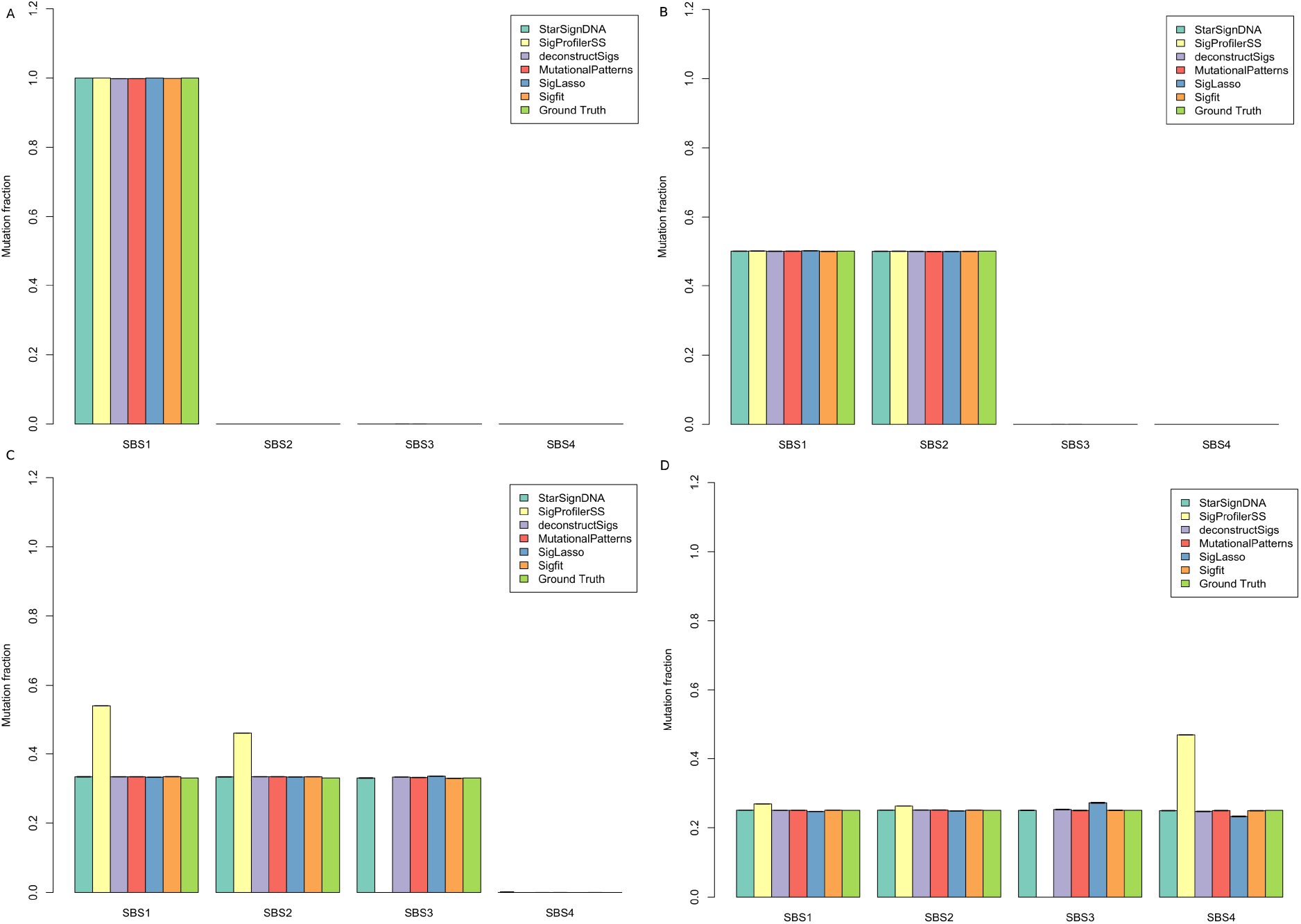
Performance comparison of six mutational signature re-fitting tools. StarSignDNA (light green), SigProfilerSS (light yellow), deconstructSigs (light blue), MutationalPatterns (red), SigLASSO (dark blue), and sigfit (orange) across four different signature types (SBS1, SBS2, SBS3, and SBS4) is represented by each set of bars. The vertical lines on top of each bar represent the average exposure of 100 simulated samples (approximately 10000 mutations each) indicating the variability or uncertainty in the mutation fraction estimation by each tool compared to the known exposure values (Ground Truth). A) Only SBS1 has a true exposure of 1, while SBS2, SBS3, and SBS4 have a true exposure of 0. B) SBS1 and SBS2 have true exposures of 0.5, while SBS3 and SBS4 have true exposures of 0. C) SBS1, SBS2, and SBS3 each have true exposures of 0.333, while SBS4 has a true exposure of 0. D) All four signatures have an equal true exposure of 0.25.

**S4 Figure.**
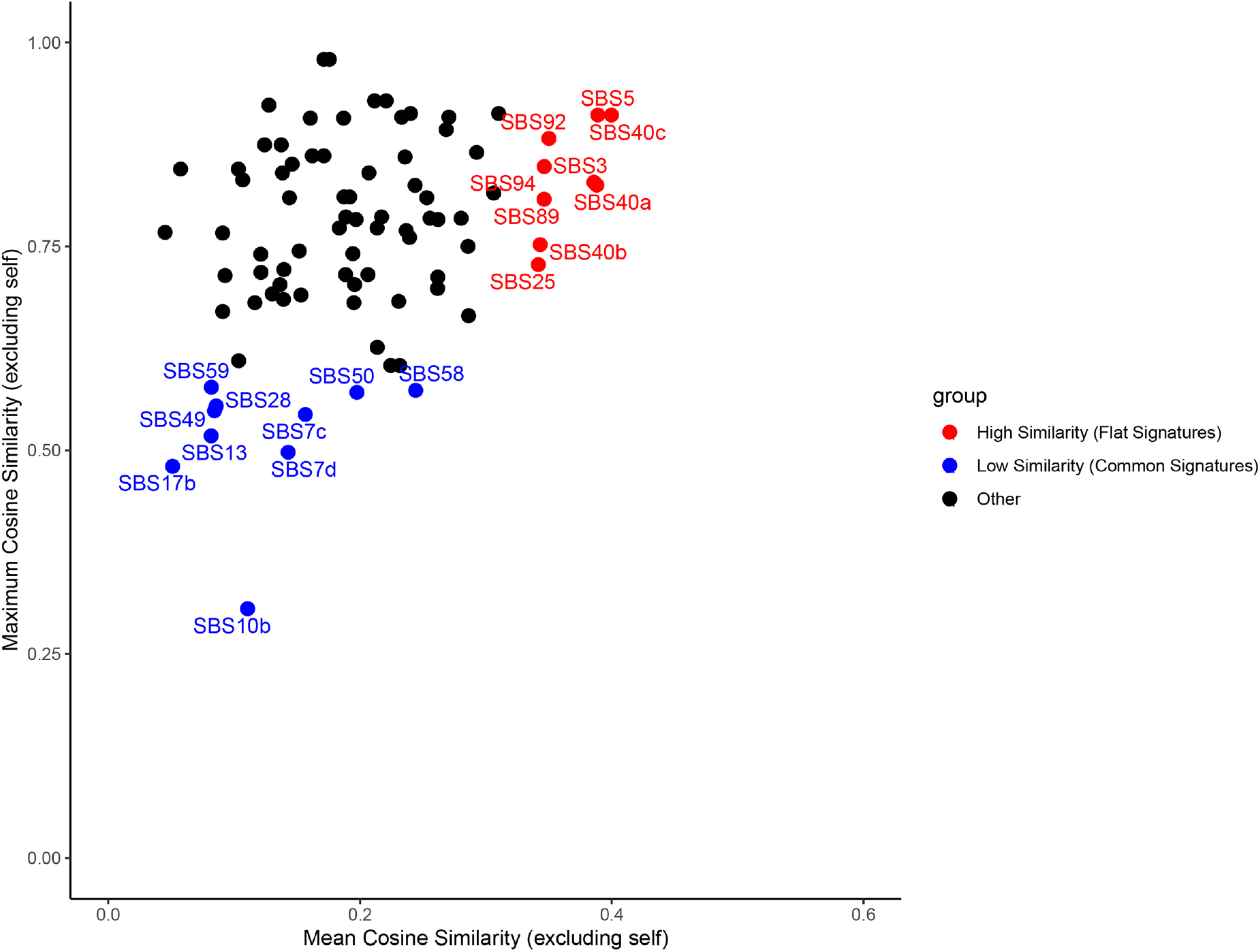
COSMIC version 3.4 reference signature Cosine Similarity distribution. The red points highlight the distribution and correlation of mutation frequencies with an average cosine similarity of 0.32 and a maximum of 0.6 for the flat signatures: SBS3, SBS5, SBS40c, SBS92, and SBS94. The blue points highlight the distribution and correlation of mutation frequencies with an average cosine similarity of 0.29 and a maximum of 0.6 for the common signatures: SBS7d, SBS10b, SBS13, SBS17b, and SBS28. The black points represent the rest of the signatures. This plot helps visualize the relationship between mean and maximum mutation frequencies for various SBS signatures.

**S5 Figure.**
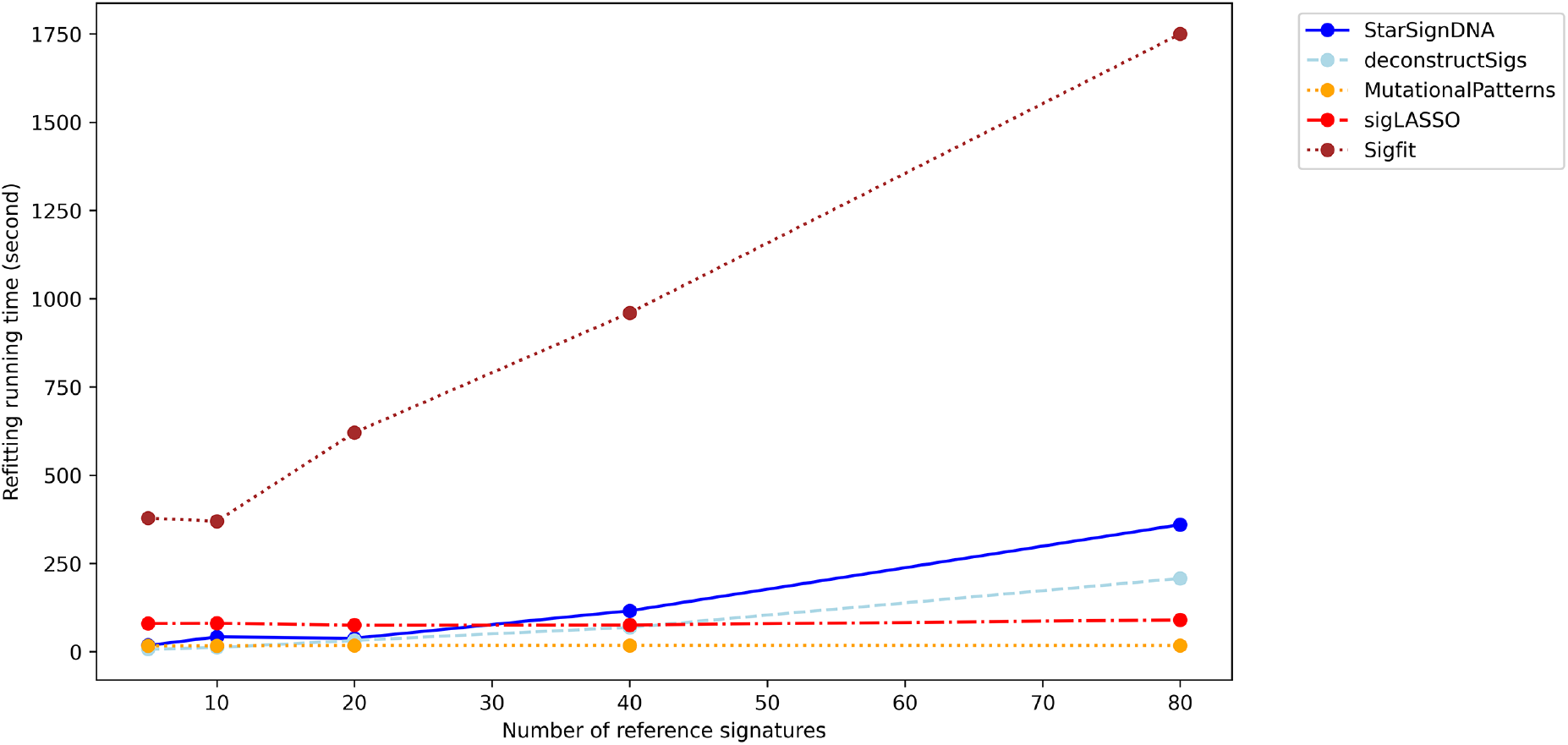
re-fitting algorithm running time. Running time of StarSignDNA, deconstructSigs, MutationalPatterns, sigLASSO, and sigfit at different numbers of signatures simulated with 100 samples of 100 mutations per sample.

**S6 Figure.**
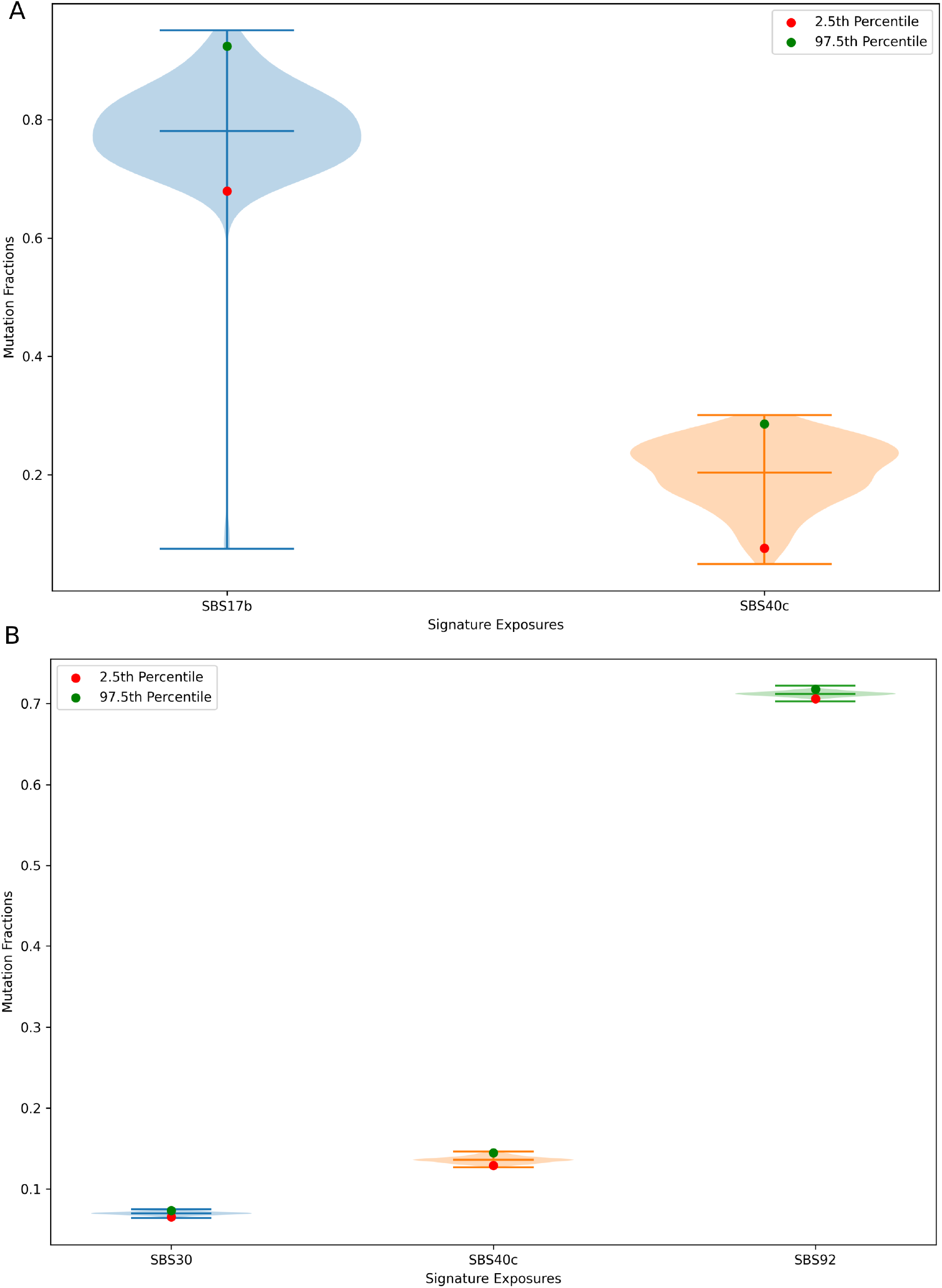
A visualization depicts the distribution of mutation fractions, showcasing the spread and concentration of the data. The y-axis represents the mutation fractions, indicating the proportion of mutations attributable to each signature. The vertical line marks the median, with colored dots highlighting the 2.5th percentile (red dot) and the 97.5th percentile (green dot). A) Represents the results from a dataset in which SBS17b was simulated. B) Represents the results from a dataset in which SBS92 was simulated.

**S7 Figure.**
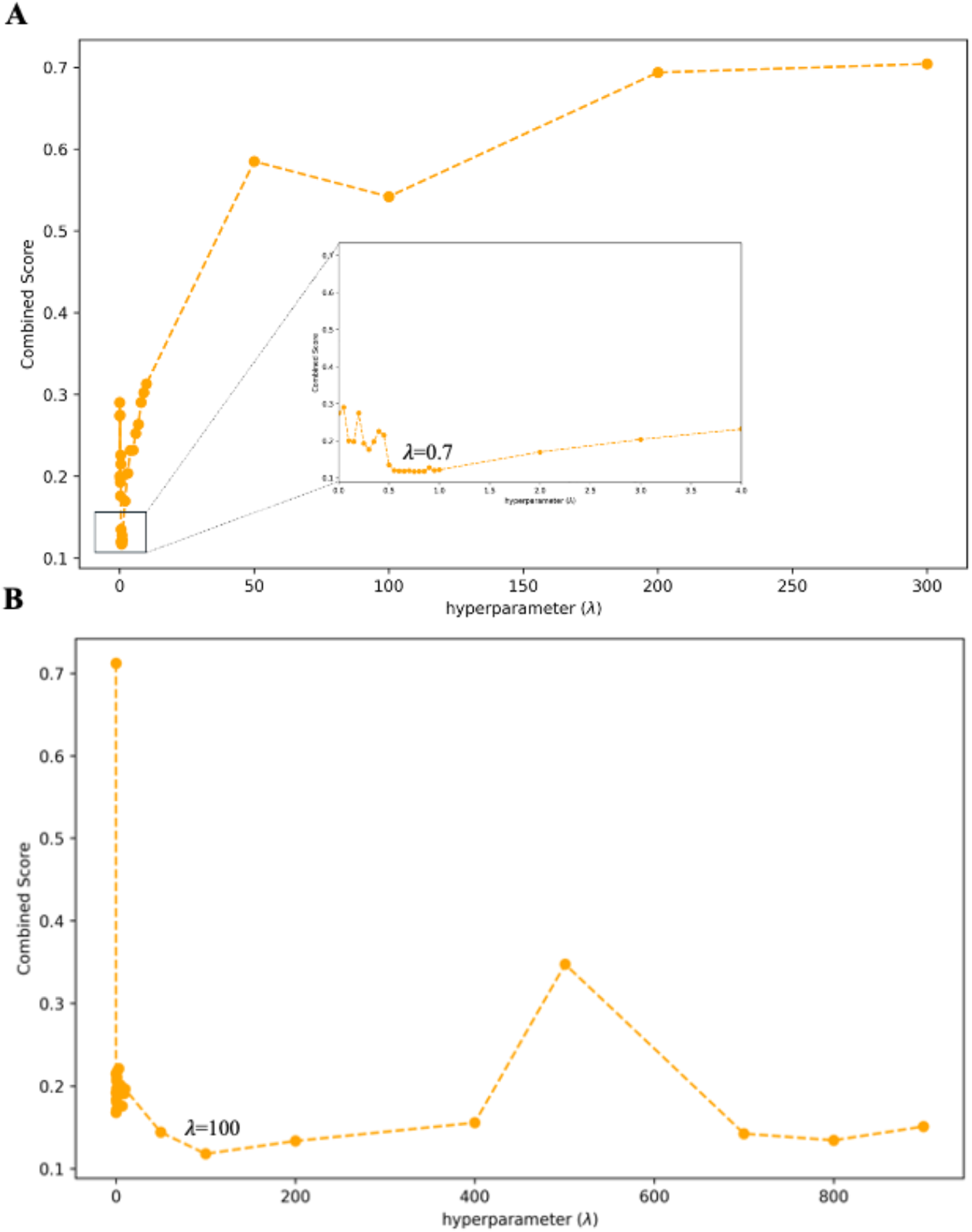
Hyperparameter *λ* obtained using cross-validation for WES (A) and WGS (B).

**S8 Figure.**
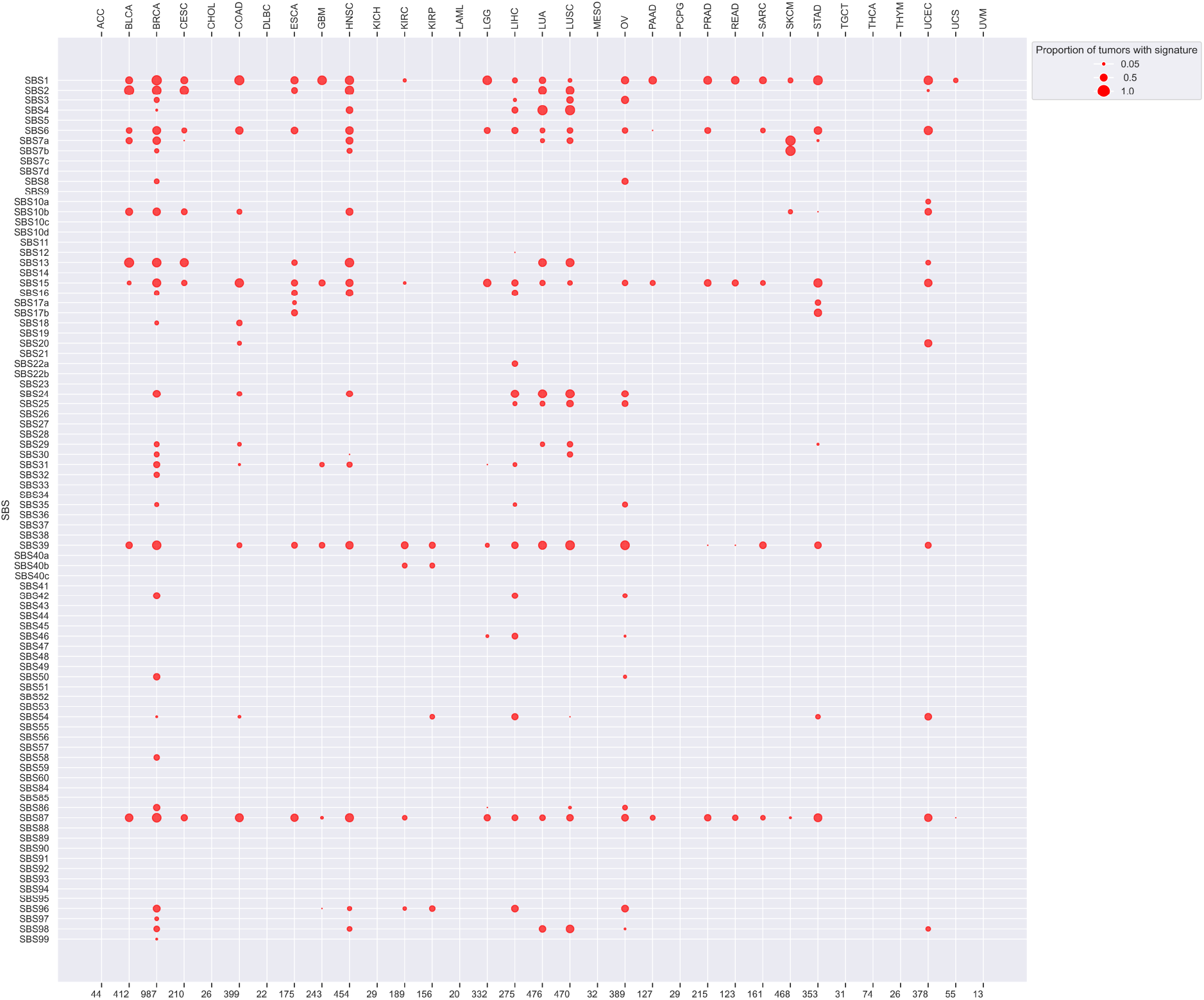
This figure shows the performance of sigLASSO on 33 TCGA cancer types listed alphabetically along the top x-axis. The size of each dot represents the proportion of signatures within a specific cancer type that exhibits a mutational signature identified by sigLASSO. The bottom x-axis indicates the number of samples analyzed for each cancer type. A separate panel legend identifies the *active signatures* (those contributing more than 6% of the mutations) in each cancer type using different methods. All 86 COSMIC v3.4 signatures were used as references for the analysis.

**S9 Figure.**
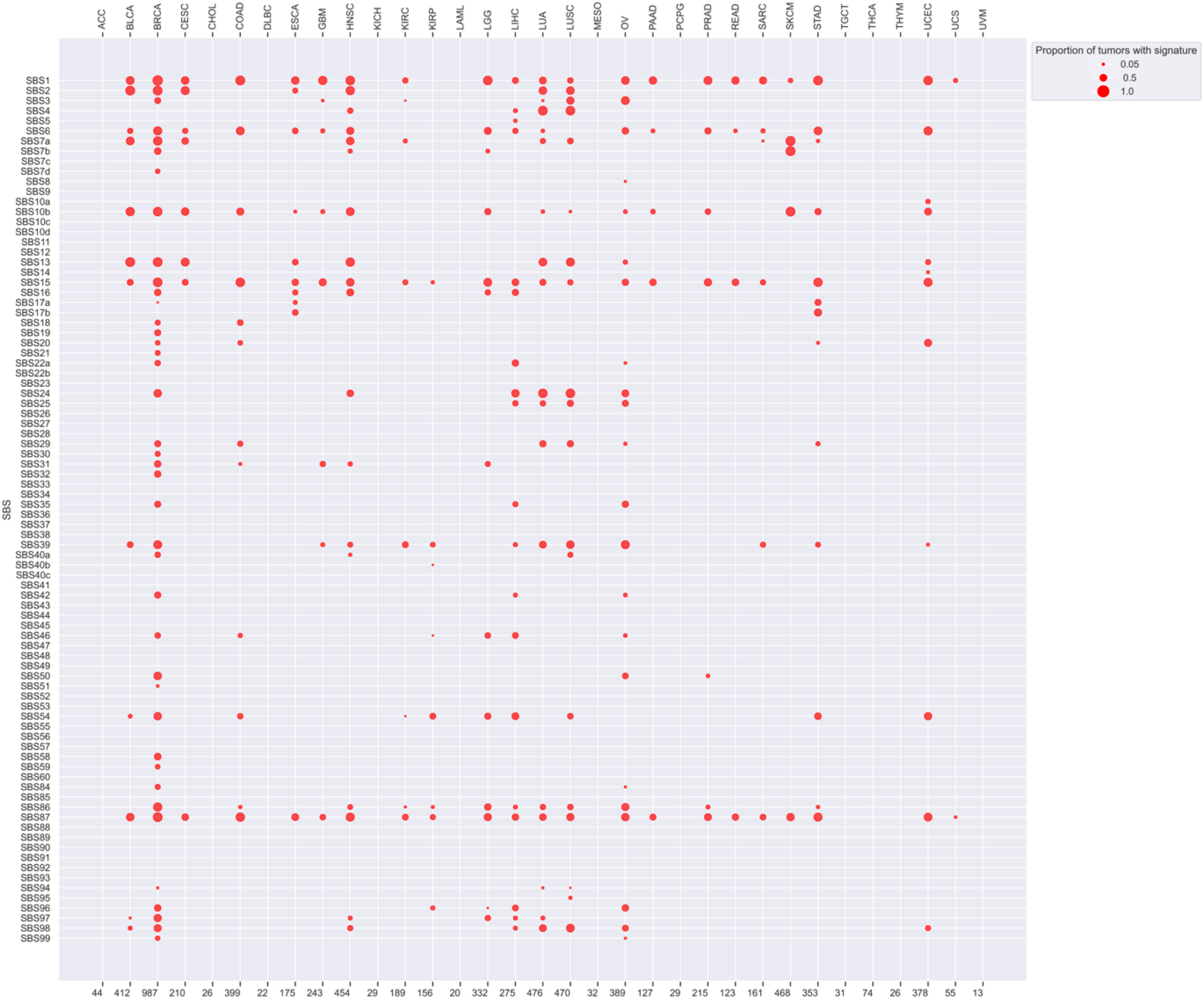
This figure shows the performance of deconstructSigs on 33 TCGA cancer types listed alphabetically along the top x-axis. The size of each dot represents the proportion of signatures within a specific cancer type that exhibits a mutational signature identified by deconstructSigs. The bottom x-axis indicates the number of samples analyzed for each cancer type. A separate panel legend identifies the *active signatures* (those contributing more than 6% of the mutations) in each cancer type using different methods. All 86 COSMIC v3.4 signatures were used as references for the analysis.

**S1 Table. Exposure matrices of six mutational signature re-fitting methods on simulated datasets**. The exposure matrix of the second category of the simulated dataset uses common signatures SBS7d, SBS10b, SBS13, SBS17b, and SBS28 as a reference signature with 100 samples, each having 100 mutations per sample.

**S2 Table. Exposure matrices of six mutational signature re-fitting methods on simulated datasets**. The exposure matrix of the second category of the simulated dataset uses common signatures SBS7d, SBS10b, SBS13, SBS17b, and SBS28 as a reference signature with 500 samples, each having 500 mutations per sample.

**S3 Table. Exposure matrices of six mutational signature re-fitting methods on simulated datasets**. The exposure matrix of the second category of the simulated dataset uses common signatures SBS7d, SBS10b, SBS13, SBS17b, and SBS28 as a reference signature with 1000 samples, each having 1000 mutations per sample.

**S4 Table. Exposure matrices of six mutational signature re-fitting methods on simulated datasets**. The exposure matrix of the second category of the simulated dataset with 100 samples, each having 100 mutations per sample. All 86 COSMIC v3.4 signatures were used as references for the analysis.

**S5 Table. Exposure matrices of six mutational signature re-fitting methods on simulated datasets**. The exposure matrix of the second category of the simulated dataset with 100 samples, each having 500 mutations per sample. All 86 COSMIC v3.4 signatures were used as references for the analysis.

**S6 Table. Exposure matrices of six mutational signature re-fitting methods on simulated datasets**. The exposure matrix of the second category of the simulated dataset with 100 samples, each having 1000 mutations per sample. All 86 COSMIC v3.4 signatures were used as references for the analysis.

**S7 Table. Exposure matrices of six mutational signature re-fitting methods on simulated datasets**. The exposure matrix of the third category of the simulated dataset uses common signatures SBS3, SBS5, SBS40c, SBS92, and SBS94 as a reference signature with 100 samples, each having 100 mutations per sample.

**S8 Table. Exposure matrices of six mutational signature re-fitting methods on simulated datasets**. The exposure matrix of the third category of the simulated dataset uses common signatures SBS3, SBS5, SBS40c, SBS92, and SBS94 as a reference signature with 500 samples, each having 100 mutations per sample.

**S9 Table. Exposure matrices of six mutational signature re-fitting methods on simulated datasets**. The exposure matrix of the third category of the simulated dataset uses common signatures SBS3, SBS5, SBS40c, SBS92, and SBS94 as a reference signature with 1000 samples, each having 100 mutations per sample.

**S10 Table. Exposure matrices of six mutational signature re-fitting methods on simulated datasets**. The exposure matrix of the third category of the simulated dataset with 100 samples, each having 100 mutations per sample. All 86 COSMIC v3.4 signatures were used as references for the analysis.

**S11 Table. Exposure matrices of six mutational signature re-fitting methods on simulated datasets**. The exposure matrix of the third category of the simulated dataset with 100 samples, each having 500 mutations per sample. All 86 COSMIC v3.4 signatures were used as references for the analysis.

**S12 Table. Exposure matrices of six mutational signature re-fitting methods on simulated datasets**. The exposure matrix of the third category of the simulated dataset with 100 samples, each having 1000 mutations per sample. All 86 COSMIC v3.4 signatures were used as references for the analysis.

**S13 Table. Exposure matrices of six mutational signature re-fitting methods on simulated datasets**. The exposure matrix of the third category of the simulated downsample dataset with 100 samples, each having 50 mutations per sample. All 86 COSMIC v3.4 signatures were used as references for the analysis.

**S14 Table. Signatures discovered using Breast cancer PCAWG data using StarSignDNA, SigneR, SigProfiler, SUITOR, and SparseSignatures algorithms**.

**S15 Table. Signatures discovered using SKIN cancer PCAWG data using StarSignDNA, SigneR, SigProfiler, SUITOR, and SparseSignatures algorithms**.

**S16 Table. Signatures discovered using Pancreatic cancer PCAWG data using StarSignDNA, SigneR, SigProfiler, SUITOR, and SparseSignatures algorithms**.

**S17 Table. Signatures discovered using Prostate cancer PCAWG data using StarSignDNA, SigneR, SigProfiler, SUITOR, and SparseSignatures algorithms**.

**S18 Table. Signatures discovered using Kidney cancer PCAWG data using StarSignDNA, SigneR, SigProfiler, SUITOR, and SparseSignatures algorithms**.

**S19 Table. Signatures discovered using Breast cancer HMF data using StarSignDNA, SigneR, SigProfiler, SUITOR, and SparseSignatures algorithms**.

**S20 Table. Signatures discovered using SKIN cancer HMF data using StarSignDNA, SigneR, SigProfiler, SUITOR, and SparseSignatures algorithms**.

**S21 Table. Signatures discovered using Pancreatic cancer HMF data using StarSignDNA, SigneR, SigProfiler, SUITOR, and SparseSignatures algorithms**.

**S22 Table. Signatures discovered using Prostate cancer HMF data using StarSignDNA, SigneR, SigProfiler, SUITOR, and SparseSignatures algorithms**.

**S23 Table. Signatures discovered using Kidney cancer HMF data using StarSignDNA, SigneR, SigProfiler, SUITOR, and SparseSignatures algorithms**.

**S24 Table. Cosine similarity obtained using Breast cancer PCAWG signatures detected versus COSMIC reference signatures for StarSignDNA, SigneR, SigProfiler, SUITOR, and SparseSignatures algorithms**.

**S25 Table. Cosine similarity obtained using Skin cancer PCAWG signatures detected versus COSMIC reference signatures for StarSignDNA, SigneR, SigProfiler, SUITOR, and SparseSignatures algorithms**.

**S26 Table. Cosine similarity obtained using Pancreatic cancer PCAWG signatures detected versus COSMIC reference signatures for StarSignDNA, SigneR, SigProfiler, SUITOR, and SparseSignatures algorithms**.

**S27 Table. Cosine similarity obtained using Prostate cancer PCAWG signatures detected versus COSMIC reference signatures for StarSignDNA, SigneR, SigProfiler, SUITOR, and SparseSignatures algorithms**.

**S28 Table. Cosine similarity obtained using Kidney cancer PCAWG signatures detected versus COSMIC reference signatures for StarSignDNA, SigneR, SigProfiler, SUITOR, and SparseSignatures algorithms**.

**S29 Table. Cosine similarity obtained using Breast cancer HMF signatures detected versus COSMIC reference signatures for StarSignDNA, SigneR, SigProfiler, SUITOR, and SparseSignatures algorithms**.

**S30 Table. Cosine similarity obtained using Skin cancer HMF signatures detected versus COSMIC reference signatures for StarSignDNA, SigneR, SigProfiler, SUITOR and SparseSignatures algorithms**.

**S31 Table. Cosine similarity obtained using Pancreatic cancer HMF signatures detected versus COSMIC reference signatures for StarSignDNA, SigneR, SigProfiler, SUITOR, and SparseSignatures algorithms**.

**S32 Table. Cosine similarity obtained using Prostate cancer HMF signatures detected versus COSMIC reference signatures for StarSignDNA, SigneR, SigProfiler, SUITOR and SparseSignatures algorithms**.

**S33 Table. Cosine similarity obtained using Kidney cancer HMF signatures detected versus COSMIC reference signatures for StarSignDNA, SigneR, SigProfiler, SUITOR, and SparseSignatures algorithms**.

## Acknowledgments

This project has received funding from the European Union’s Horizon 2020 research and innovation program under the Marie Sk-lodowska-Curie grant agreement No 801133.

## Author Contributions

### Conceptualization

Christian Domilongo Bope, Sigve Nakken, Ole Christian Lingjærde, and Eivind Hovig.

### Developed the mathematical framework

Christian Domilongo Bope and Ole Christian Lingjærde.

### Software

Christian Domilongo Bope, Knut D. Rand and Sumana Kalyanasundaram.

### Performed the simulations, benchmarking, and results validation

Christian Domilongo Bope and Sumana Kalyanasundaram.

### Interpretation of results and manuscript preparation

Christian Domilongo Bope, Sumana Kalyanasundaram, Sigve Nakken, Ole Christian Lingjærde and Eivind Hovig.

**All the authors critically reviewed and approved the final manuscript**.

## StarSignDNA Supporting Document

### Simulations datasets

To create the fifth category of simulated datasets, we followed these steps:

1. Data Acquisition: We obtained 146 whole-genome sequencing (WGS) datasets from prostate cancer samples. These datasets were retrieved from the PCAWG project hosted by the International Cancer Genome Consortium (ICGC) (https://dcc.icgc.org/pcawg).
2. Signature Selection: We selected a set of four underlying mutational signatures from COSMIC version 3.4. These signatures were SBS1, SBS5, SBS18, and SBS40, which are known to be active processes specifically relevant to prostate cancer [**1**].
3. Signature Deconvolution: We employed the DeconstructSigs algorithm [**2**] to fit the chosen signatures to the WGS data. This process reconstructed a mutational catalog count, denoted as *M*\* as follows:
  - Let **M** *∈* ℝ^*N*,96^ be a mutational signature catalogue;
  - **S** = (*s*_*jk*_) *∈* ℝ^96*×K*^ where *K* is four, representing COSMIC signatures SBS1, SBS5, SBS18 and SBS40
  - The exposure matrix **E** was estimated using deconstructSigs refitting algorithm
  - The 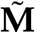 mutational catalogue over-weighted matrix on signatures SBS1, SBS5, SBS18 and SBS40 was obtained as follows:
  - 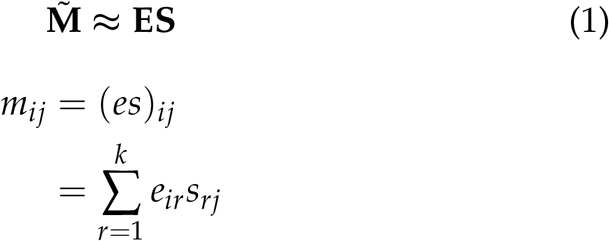

where *m*_*ij*_ is an element of 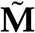.
4. We generated 50 bootstraps without replacement using [**3**].

### Tuning hyperparameter *λ*

The level of regularization of the mutational exposures obtained by LASSO is defined by *λ*. For the refitting algorithm, *λ* is obtained by computing using cross-validation. The optimal parameter is obtained by computing the combined score. Let *W*_*mse*_ be the weight of the Mean Square Error (0.2 and 0.1 for WES and WGS, respectively); and *W*_*sparse*_ be the weight of sparsity (0.1 and 0.7 for WES and WGS, respectively), and let *W*_*pm f*_ be the weight of the probability mass function (PMF) (0.7 and 0.2 for WES and WGS, respectively). We first normalize MSE, Sparsity, and PMF.

- Normalized MSE 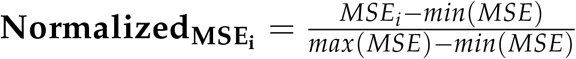
- Normalized Sparsity 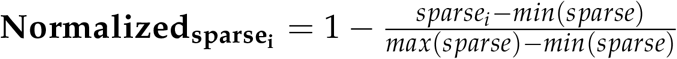
- Normalized PMF 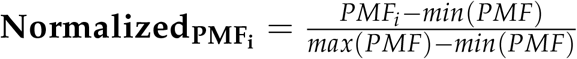

Then, we compute the combined score as follows:

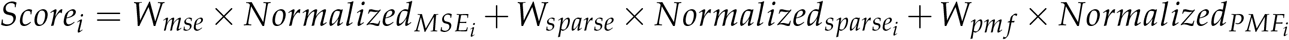

The hyperparameters correspond to the smallest combined score.

### Parameters used for other mutational signatures analysis tools

The parameters used for other mutational signature analysis tools for refitting and *de novo* are listed as follows:

#### Re-fitting tools

The default parameters for each of the programs were used for SigLASSO, deconstructSigs, MutationalPatterns, Sigfit, and SigProfiler Single Sample.

#### *De novo* tools

For the *de novo* tools, the following settings were applied:

**SigneR:** signeR(M=t(input), nlim=c(2,18), try all=TRUE)

**SUITOR:** minimum rank: 2, maximum rank: 18; number of folds:10; EM algorithm stopping tolerance: 1*e* − 5, maximum number of iterations in EM algorithm: 2000; number of seeds: 10000

**SigProfilerExtractor:** sig.sigProfilerExtractor(“matrix”, “output folder name”, data, minimum signatures=2, maximum signatures=18, nmf replicates=100, cpu=-1)

**SparseSignatures:** Number of signatures: 2 to 18 *λ* (sparsity): 0.01,0.02,0.05 and 0.1

Number of repetitions of NMF to calculate initial values: default

Number of iterations to fit signatures using the alternating method with sparsity: 30

Number of restarts per repetition of cross-validation: 5 Number of repetitions of bi-cross-validation: default

Background signature: Default

